# Disrupted memory T cell expansion in HIV-exposed uninfected infants is preceded by premature skewing of T cell receptor clonality

**DOI:** 10.1101/2023.05.19.540713

**Authors:** Sonwabile Dzanibe, Aaron J. Wilk, Susan Canny, Thanmayi Ranganath, Berenice Alinde, Florian Rubelt, Huang Huang, Mark M. Davis, Susan Holmes, Heather B. Jaspan, Catherine A. Blish, Clive M. Gray

**Affiliations:** Division of Immunology, Department of Pathology, Institute of Infectious Disease and Molecular Medicine, University of Cape Town, Cape Town, South Africa; Department of Medicine, School of Medicine, Stanford University, Stanford, CA; Division of Rheumatology, Department of Pediatrics, Seattle Children’s Hospital, Seattle, WA USA; Division of Molecular Biology and Human Genetics, Stellenbosch University, Cape Town, South Africa; Department of Microbiology and Immunology, Stanford University School of Medicine, Stanford, CA, USA; Institute for Immunity, Transplantation and Infection, Stanford University School of Medicine, Stanford, CA, USA; Howard Hughes Medical Institute, School of Medicine, Stanford University, Stanford, CA; Department of Statistics, Stanford University, Stanford, CA, USA; Seattle Children’s Research Institute and Department of Paediatrics and Global Health, University of Washington, Seattle, WA; Chan Zuckerberg Biohub, San Francisco, CA

**Keywords:** HIV exposure, infant immunity, T cell receptor, NK cels and antibody responses

## Abstract

While preventing vertical HIV transmission has been very successful, the increasing number of HIV-exposed uninfected infants (iHEU) experience an elevated risk to infections compared to HIV-unexposed and uninfected infants (iHUU). Immune developmental differences between iHEU and iHUU remains poorly understood and here we present a longitudinal multimodal analysis of infant immune ontogeny that highlights the impact of HIV/ARV exposure. Using mass cytometry, we show alterations and differences in the emergence of NK cell populations and T cell memory differentiation between iHEU and iHUU. Specific NK cells observed at birth were also predictive of acellular pertussis and rotavirus vaccine-induced IgG and IgA responses, respectively, at 3 and 9 months of life. T cell receptor Vβ clonotypic diversity was significantly and persistently lower in iHEU preceding the expansion of T cell memory. Our findings show that HIV/ARV exposure disrupts innate and adaptive immunity from birth which may underlie relative vulnerability to infections.

## Introduction

Early life, especially in Africa, is often plagued by high infectious morbidity with infectious diseases accounting for 2.61 million deaths of which 45% occur duing the neonatal period^1^. This high mortality rate is most likely linked to the period during which the immune system adapts to extrauterine life. During this transition window, several factors such as maternal morbidity, microbial exposure, and vaccination significantly modulate infant immunity and consequently influences health and disease outcomes^2–5^. Investigating immune ontogeny and factors that impact immune trajectory during infancy can provide important insights into understanding how better to combat these early life immune stressors.

Current dogma is that the *in utero* environment is sterile and that newborn infants have limited antigen exposure prior to birth, with T cells being predominantly naïve with little T cell receptor (TCR) engagement^6, 7^. This lack of pre-existing adaptive cellular memory early in life is likely to increase the vulnerability of infants to infectious agents and disease, especially if there is an absence of adequate breast-feeding to provide passive maternal antibody immunity^3, 8, 9^. Since the diversity and composition of the TCR is presumably dependent on prior antigen exposure, examining changes in the TCR repertoire during infancy could provide insight into immune development and how specific immune modulatory factors influence susceptibility to infectious diseases.

The advent of the Option B+ vertical transmission prevention program with effective antiretroviral (ARV) drugs has ensured that HIV transmission has been minimised, but also introduces the possible adverse effect of HIV and ARV exposure on neonatal development *in utero* and in the post-natal period. The intertwined nature of HIV and ARV exposure makes it difficult to unravel, but it is known that HIV/ARV-exposed uninfected infants (iHEU) have higher risk of infectious disease-related morbidity and mortality compared to HIV-unexposed uninfected infants (iHUU) of a similar age, suggesting disruptions to their immune maturation^10–15^. Many studies investigating immunological disparities between iHEU and iHUU are limited to cross-sectional analysis with few studies investigating longitudinal changes and thus there is a lack of evidence on how T cells mature early in life in the context of HIV/ARV exposure. We have previously shown that maternal HIV/ARV exposure alters the dynamics of the T regulatory (Treg) to Th17 cell ratio resulting in a Th17/Treg imbalance associated with gut damage^16^. In this paper, we extend this analysis to investigate HIV/ARV exposure on T cell clonality, memory and NK cell maturation differences between iHEU and iHUU.

Since early life is marked by inexperienced adaptive immunity, neonatal immune defence is heavily reliant on innate immune cells such as NK cells to rapidly eliminate infections. Compared to adults, however, neonatal NK cells display functional defects such as reduced cytolytic activity including antibody mediated cell cytotoxicity^17, 18^, decreased expression of adhesion molecules^19^ and lower secretion of TNF-α and INF-γ^20^. Earlier studies have further demonstrated that compared to iHUU, iHEU have a lower proportion of NK cells measured at birth and 6 months, and having reduced IFN-γ secretion and perforin expression^21, 22^. In addition, the function of NK cells is influenced by the combinatorial signalling through a diverse array of activating and inhibitory receptors expressed on the cell surface^23^. The composition of these receptors is driven by past viral exposure^20, 24, 25^. Given that NK cells also play a key role in priming the adaptive immune system to respond to invading pathogens or to tailor their immunogenicity towards specific vaccines^26^, it is important to investigate the maturation trajectory of NK cells and how this could shape adaptive immunity. Whether early life exposure to HIV/ARV shapes the maturation and/or diversification of NK cells remains unknown.

Here, we comprehensively investigated the relationship between adaptive and innate immunophenotypes along with TCR diversity and the ability to mount an antibody response to pertussis and rotavirus vaccination. Given the heightened infectious morbidity risk in iHEU, we hypothesised that immune ontogeny is associated with a narrowing TCR repertoire over time in parallel with expanded NK cell subsets in iHEU relative to iHUU. We mapped the immune trajectory of NK and T cell phenotypic clusters in the first 9 months of life and associated this with TCR diversity and antibody titres. Using this approach, we show divergent adaptive immune maturation in iHEU relative to iHUU occurring from 3 months of life with earlier phenotypic differences evident in NK cells. Our data show a narrowing and persistent skewing of the TCR repertoire from birth in iHEU that precedes altered CD4^+^ and CD8^+^ T cell memory development.

## Results

### Immune cell transition from birth to 9 months of age

We first analysed the immune ontogeny of innate and adaptive immune cell subsets and marker expression profiles longitudinally during the first 9 months in our iHUU and iHEU samples. Infant samples were analysed longitudinally using a mass cytometry antibody panel (Table S1). This included matched PBMC at birth (n=44) and at week 4 (n=53), 15 (n=52) and 36 (n=53), as well as in some cord blood mononuclear cells (n=5). Unsupervised cell clustering on all samples using the live singlet cell population (Figure S1A) revealed 11 cell clusters, shown in Figure 1A, consisting of B cells (cluster 1), T cells (CD4^+^ T cluster 2 and CD8^+^ T clusters 3, 4, and 9), monocytes (cluster 8), NK cells (cluster 10), and NKT-like cell (cluster 11). The three CD8^+^ T cell clusters (3, 4 and 9) were delineated as naïve (13.9%, CD45RA^+^CD27^+^CCR7^+^), a small effector memory population (0.07%, EM: CD45RA^+^CD27^+^CCR7^-^) and effector cells (4.4%, Eff: CD45RA^+^CD27^-^CCR7^-^). We also identified a lineage negative cluster (3.2%) and one cluster expressing HLA-DR only (1.2%) that appeared most closely related to the monocyte cluster (Figure 1A & S1B). Although we employed computational doublet removal from our clustering analysis, a minor miscellaneous population (0.65%) co-expressing typically mutually exclusive lineage markers was identified (Figure 1A & S1B). This small population is shown as a central cluster 7 in the dimensional reduction of the immune cell clusters (Figure 1A) and was excluded from downstream analysis.

**Figure 1:**
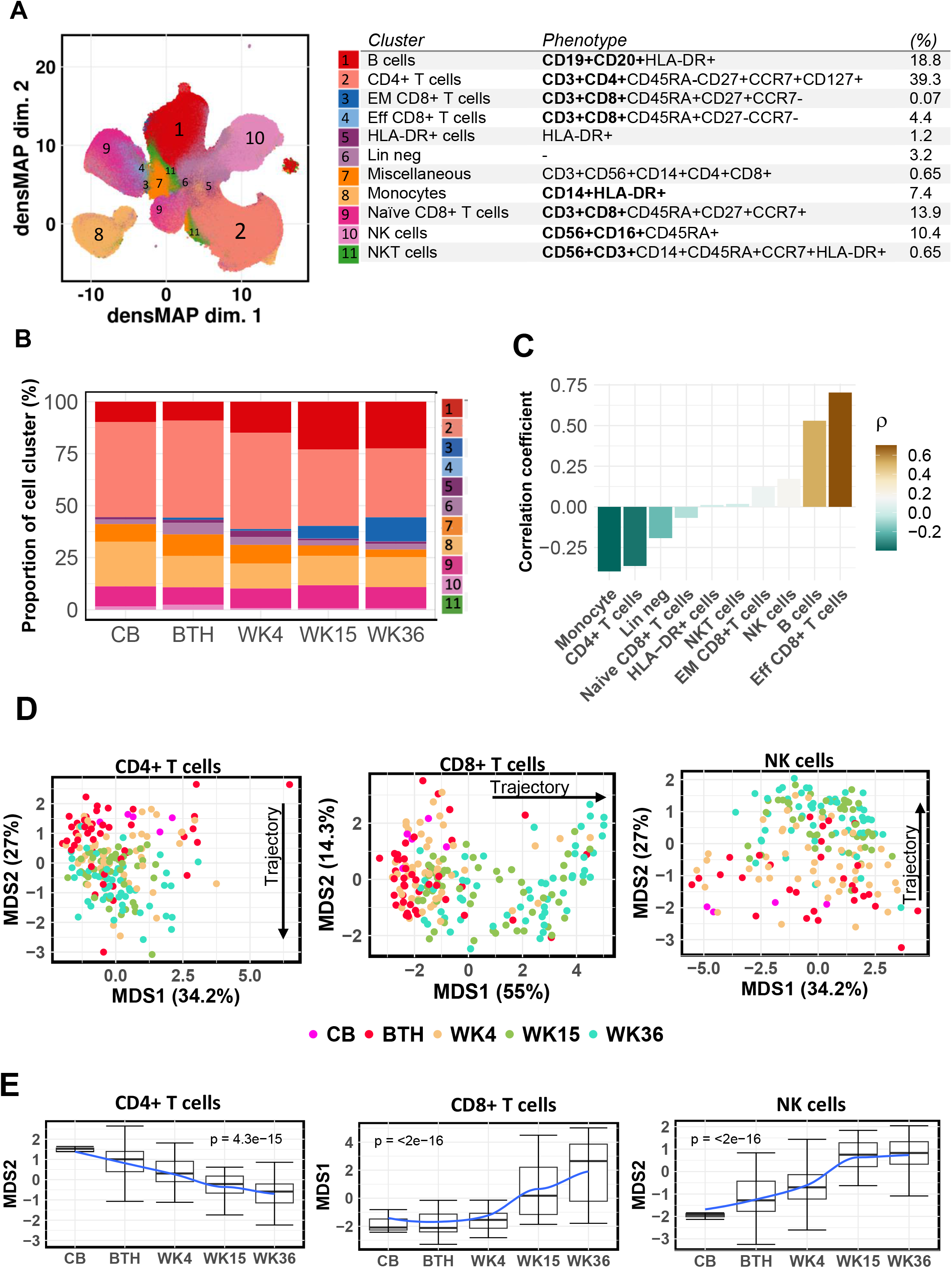
Immunophenotypic trajectory of infant peripheral blood mononuclear cells (PBMC) in the first 9 months of life. **A)** Dimensional reduction of infant PBMC Immune lineage clusters embedded on uniform manifold approximation and proximation (UMAP) with density preservation and table summarising immune cell cluster phenotype. **B)** Relative abundance of immune lineage clusters measured in cord blood (CB) and infant peripheral blood at birth (BTH), weeks (WK) 4, 15 and 36. **C)** Spearman’s rank correlation between proportion of immune lineage cell clusters and infants age from birth until week 36. **D)** Age related maturation trajectory of CD4+ and CD8+ T cells and NK cells depicted using multidimensional scaling coordinates derived from median marker expressions for each infant samples. **E)** Boxplots summarizing MDS coordinate associated with infant age for CD4+ and CD8+ T cells and NK cells.

When we examined these clusters over time, we observed an age dependent segregation of cells. Proportions of clusters changed over 36 weeks of life, where monocytes and CD4+ T cells diminished with age and B cells and effector memory CD8+ T cells increased (Figure 1B). Monocyte (*ρ=-*0.4, p <0.001) and CD4^+^ T cell (*ρ=-* 0.36, p <0.001) clusters were negatively correlated with infant age, whereas the EM CD8^+^ T cell and B cell clusters were positively correlated (*ρ=*0.66 and *ρ=*0.54 respectively, p value <0.001; Figure 1C). We further manually gated on CD4^+^ and CD8^+^ T cells and targeted the NK cell population using lineage-exclusion manual gating (Figure S1A). Using the median marker expression for each sample, multidimensional scaling (MDS) revealed intra-individual variability associated with infant age (Figure 1D). For both CD4^+^ T cells and NK cells, the grouping of the samples by marker expression displayed a converging trajectory along the MDS2-axis (Figure 1D). Both of these cell populations had a linear converging trajectory, although stabilizing from week 15 for NK cells (Figure 1E). CD8^+^ T cells, however, revealed a divergent trajectory along the MDS1-axis (Figure 1D), with a significant expansion by week 15 (Figure 1E).

Overall, these detailed unsupervised phenotypic changes in cell clustering illustrate the archetypal progression and transition of infant immunity from neonatal to an adult-like immune phenotype by 9 months of age, being characterised by a shift from innate myeloid cells to an increase in lymphoid adaptive immune cells.

### CD4 and CD8 T cell memory maturation is disrupted by HIV exposure after three months of life

To identify in more depth how HIV/ARV exposure disrupts T cell phenotypes, we parsed manually gated CD4 and CD8 cells from all samples and time points using unsupervised cell clustering which generated 13 CD4^+^ T cell clusters and 7 CD8^+^ T cell clusters (Figures 2A and 2D). Tables 1 & 2 ascribe the phenotype for each of the CD4^+^ and CD8^+^ T cell clusters, respectively. The marker expression patterns and relationships between the cell clusters are shown using unsupervised hierarchical clustering heatmaps in Figure S2A.

**Figure 2:**
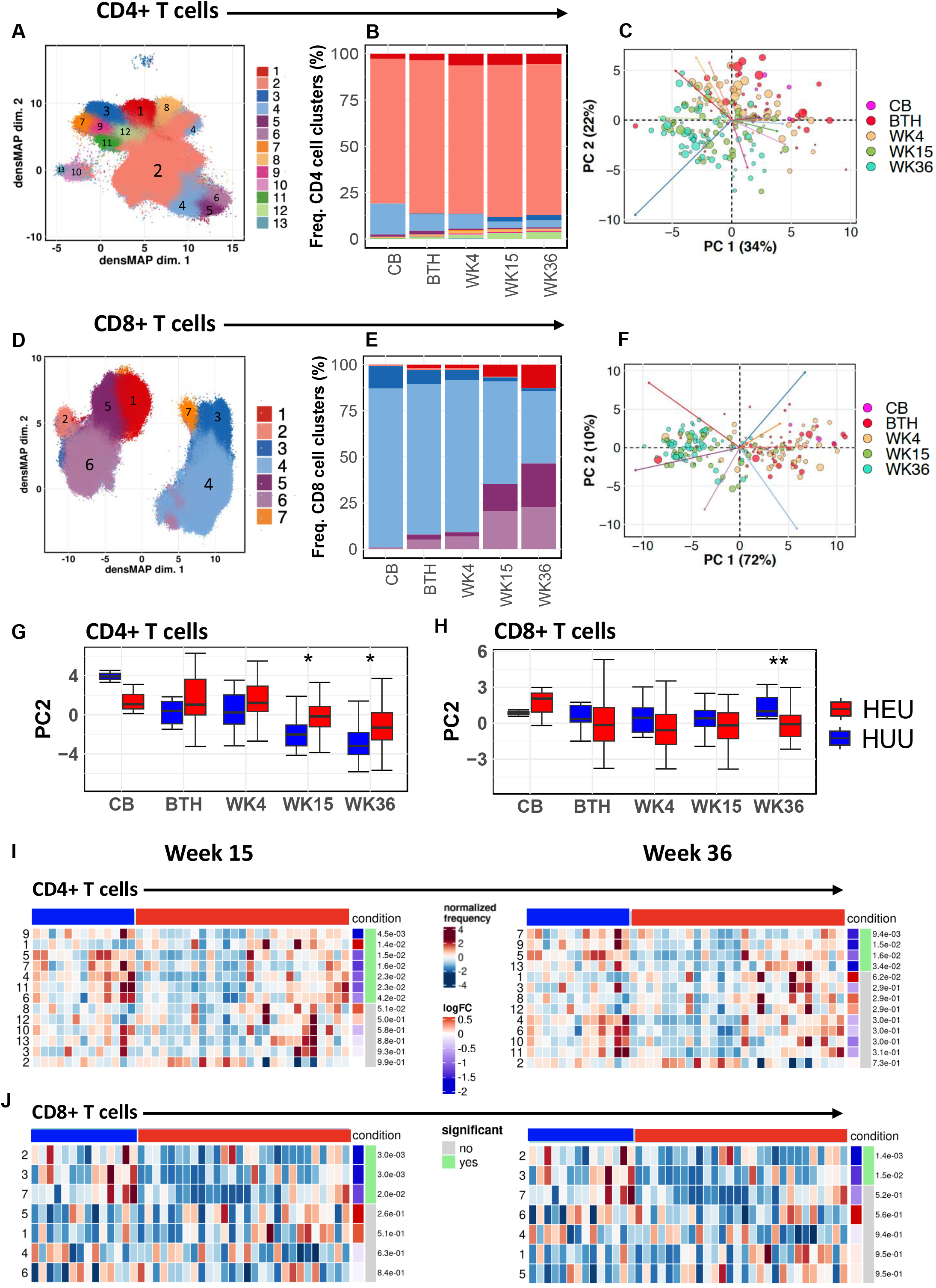
Divergent T cell memory differentiation in HIV-exposed uninfected infants (iHEU) compared to HIV-unexposed uninfected infants (iHUU). **A & D)** Uniform manifold approximation and proximation (UMAP) with density preservation showing dimensional reduction of FlowSOM clusters of CD4+ and CD8 T cells **B &E)** Relative abundances of CD4+ and CD8 + T cell clusters measured in cord blood (CB) and infant peripheral blood at birth (BTH), weeks (WK) 4, 15 and 36. **C &F)** Age related phenotypic composition of CD4+ and CD8+ T cells depicted as principal component (PC) coordinates of centred log-odd ratios of FlowSOM clusters for each infant samples and arrows indicating contribution of each cell cluster in scatter of PC components. **G &H)** Boxplots comparing PC coordinates of CD4+ and CD8+ T cells between iHEU and iHUU longitudinally. **I & J)** Generalized linear mixed model (GLMM) comparing the abundances of CD4+ and CD8+ T cell clusters between iHEU and iHUU at weeks 15 and 36.

Across all time points, 13 clusters of CD4+ T cells existed (Figure 2A and Table 1) where naïve cells (cluster 2, CD45RA^+^CD27^+^CCR7^+^) made up the predominant population with an overall frequency of 84.1% (Figure 2A & Table 1). The remaining 12 clusters for CD4+ T cells consisted of minor populations with the top three being activated naïve-like cells (cluster 4, CD38^+^HLA-DR**^+^**CD45RA^+^CD27^+^CCR7^+^), PD-1 expressing activated cells (cluster 12, PD-1+CD38+) and cytotoxic terminally differentiated cells (cluster 3, CD45RA^+^CD27^-^CCR7^-^CD57^+^Per^+^PD-1^+^). When these cell clusters were examined over infant age, there was a proportional increase in cluster 1 (Th2-like, CCR4^+^CCR6^-^CXCR3^-^), decrease in cluster 4 (activated cells, CD38^+^HLA-DR^+^CD45RA^+^CD27^+^CCR7^+^) and maintenance of cluster 2 (naïve, CD45RA^+^CD27^+^CCR7^+^) (Figure 2B). We also assessed these compositional differences by performing Principal Component Analysis (PCA) on centred log-ratios of the relative cluster abundance for each sample. Consistent with the observed immunological trajectory shown in Figures 1D, phenotypic compositions were strongly influenced by infant age, where early time points (cord, birth, and week 4) were grouped together and distinguishable from week 15 and 36 time points (Figure 2C). Looking at CD8^+^ T cells in the same manner, we identified 7 cell clusters, being less diverse than CD4^+^ T cells, but a more distinct clustering between naïve and effector/terminally differentiated memory (Figure 2D and Table 2). It was also clear that naïve CD8^+^ T cells were less populous and that clusters 1 (terminally differentiated, Perforin^+^CD45RA^-^CD27^-^CCR7^-^CD57^+^), 5 (cytotoxic cells, Perforin^+^CD57^+^) and 6 (cytotoxic effector memory, Perforin^+^CD45RA^-^CD27^-^CCR7^-^) made up a higher proportion of cells than in CD4^+^ T cell compartment. When separating out these clusters over time, these perforin-expressing cell clusters (1, 5 and 6) increased over 9 months of life (Figure 2E), with a contraction of the naïve cell population. Like CD4^+^ T cells, there was an age dependent CD8^+^ T cell compositional change (Figure 2F).

**Table 1:** Phenotypic description of CD4^+^ T cell clusters idenfied by unsupervised cell clustering.

| Cluster | Phenotype | (%) |
| --- | --- | --- |
| (1) | CCR4+CCR6-CXCR3- (Th2) | 5.07 |
| (2) | CD45RA+CD27+CCR7+ (Naïve) | 84.1 |
| (3) | CD45RA+CD27-CCR7-CD57+Per+PD-1+ (Cytotoxic TD) | 1.3 |
| (4) | CD38+HLA-DR+CD45RA+CD27+CCR7+ (Activated cells) | 4.7 |
| (5) | CCR4+CCR6+CXCR3+CD57- | 0.7 |
| (6) | CCR4+CCR6+CXCR3+CD57+ | 0.2 |
| (7) | Per+Ki67+ | 0.3 |
| (8) | CD127-CD25+CD39+TIGIT+ (Treg cells) | 0.9 |
| (9) | Per+Ki67+CD25+ | 0.3 |
| (10) | Per+Ki67-HLA-DR+CD69+CD39+ | 0.1 |
| (11) | Per+Ki67+CD27+FcεRIγ+ | 0.2 |
| (12) | PD-1+CD38+ | 2.07 |
| (13) | CD39+CD45RA+CCR7-CD27- | 0.1 |

**Table 2:** Phenotypic description of CD8^+^ T cell clusters identified by unsupervised cell clustering.

| Cluster | Phenotype | (%) |
| --- | --- | --- |
| (1) | Per+CD45RA-CD27-CCR7-CD57+ (TD) | 7.5 |
| (2) | CD56+CD16+NkpK6+NKG2A+ (NKT-like) | 0.4 |
| (3) | HLA-DR+ | 2.7 |
| (4) | CD45RA+CD27+CCR7+ (Naïve) | 55.3 |
| (5) | Per+CD57+ | 15.6 |
| (6) | Per+CD45RA-CD27-CCR7- | 18.4 |
| (7) | Ki67+CD45RA+CD27+ | 0.1 |

We next determined whether there were any disruptions in T cell cluster compositions due to HIV exposure. There were statistically significant differences in PCA-2 at weeks 15 and 36 for CD4^+^ T cells between iHEU and iHUU (Figure 2G) and a converse difference at week 36 for CD8^+^ T cells (Figure 2H). To account for these divergent T cell phenotypes, we used differential abundance testing to determine which of the cell clusters were responsible for these differences. CD4 clusters 4, 5, 6, 7, 9 and 11 were significantly lower in iHEU compared to iHUU at week 15, with clusters 5, 7 and 9 remaining lower in iHEU at week 36 (Figure 2I). These less frequent of cell clusters in iHEU were characterised as either being activated (cluster 4: CD38^+^HLA-DR^+^), expressing perforin (clusters 7, 9 and 11) or exhibiting terminal differentiation (CD57^+^/PD-1^+^; clusters 5 and 6). In contrast, the CD4^+^ Th2-like cluster 1 was significantly higher in iHEU compared to iHUU at week 15 but was no longer significantly elevated by week 36 (Figure 2I). Moreover, CD4^+^ Th2 cells (cluster 1) and cytotoxic terminally differentiated cells (cluster 3: CD45RA^+^CD27^-^CCR7^-^ CD57^+^Perforin^+^PD-1^+^) were also higher in iHEU compared to iHUU at week 4 (p=0.007 and p=0.013, respectively) although these differences were not statistically significant following FDR correction (Figure S2B).

For CD8^+^ T cells, there were significantly lower mean frequencies for clusters 2 (NKT-like cells: CD56^+^CD16^+^NKp30^+^NKp46^+^NKG2A^+^Per^+^), 3 (Naïve-like activated cell: CD45RA^+^CD27^+^CCR7^+^CXCR3^+^CCR4^+^HLA-DR^+^) and 7 (proliferating EM: Ki67^+^ CD45RA^+^CD27^+^CCR7^-^Per^+^) in iHEU compared to iHUU at week 15 (Figure 2I). By week 36, only the mean frequencies of clusters 2 (NKT-like cells) and 3 (Naïve-like cells) remained significantly lower in iHEU compared to iHUU (Figure 2J).

A similar analytical approach was undertaken for NK cells, and of the 10 clusters identified (Figure 3A), 9 were CD56^dim/+^CD16^dim/+^ (Table 3). This included CD56^Hi^CD16^-^ (cluster 9), which had high NKG2A expression, six clusters that were CD56^dim/+^CD16^dim/+^ (clusters 1, 4, 5, 6, 8 and 10) that were predominantly expressing perforin, and 2 clusters that were CD56^-^CD16^+^ (clusters 3 and 7). Only cluster 2 was characterised as CD56^-^CD16^-^ and expressed DNAM1 (Figure S2A & Table S6). The NK cell phenotypic composition was also observed to be strongly associated with infant age (Figure 3B and C), with cluster 1 (Per^+^CD45RA^+^Fc_ɛ_RIIγ^+^) expanding over 36 weeks and cluster 2 (CD56^-^CD16^_^DNAM1^+^) contracting (Figure 3B). Of note, the perforin-expressing cluster 5 was expanded at weeks 15 and 36. Compositional differences in NK cell clusters between iHEU and iHUU were observed at week 4, earlier than for T cells, and while this trend persisted at week 15, the differences were not statistically significant (Figure 3D). Differential abundance testing revealed that at week 4, iHEU had higher frequencies of cluster 1 (CD56^dim^CD16^dim^) co-expressing Per^+^CD45RA^+^FcERIy+ (p=0.04, p.adj=0.2) and cluster 5 (CD56^dim^CD16^+^) co-expressing Per^+^CD57^+^CD45RA^+^CD38^+^ (p=0.03, p.adj=0.2) compared to iHEU (Figure S2C). This trend continued at week 15 for cluster 5 (p=0.041, p.adj=0.20). Also at week 15, we observed lower frequencies of cluster 10 (CD56^+^CD16^+^: p=0.027, p.adj=0.2) in iHEU compared to iHUU (Figure S2C). These findings show that that the ontogeny of immune cells is ordered, and that HIV/ARV exposure sequentially disrupts immune trajectory over time after birth beginning with that of innate (NK) cells and later disrupting memory T cell maturation.

**Figure 3:**
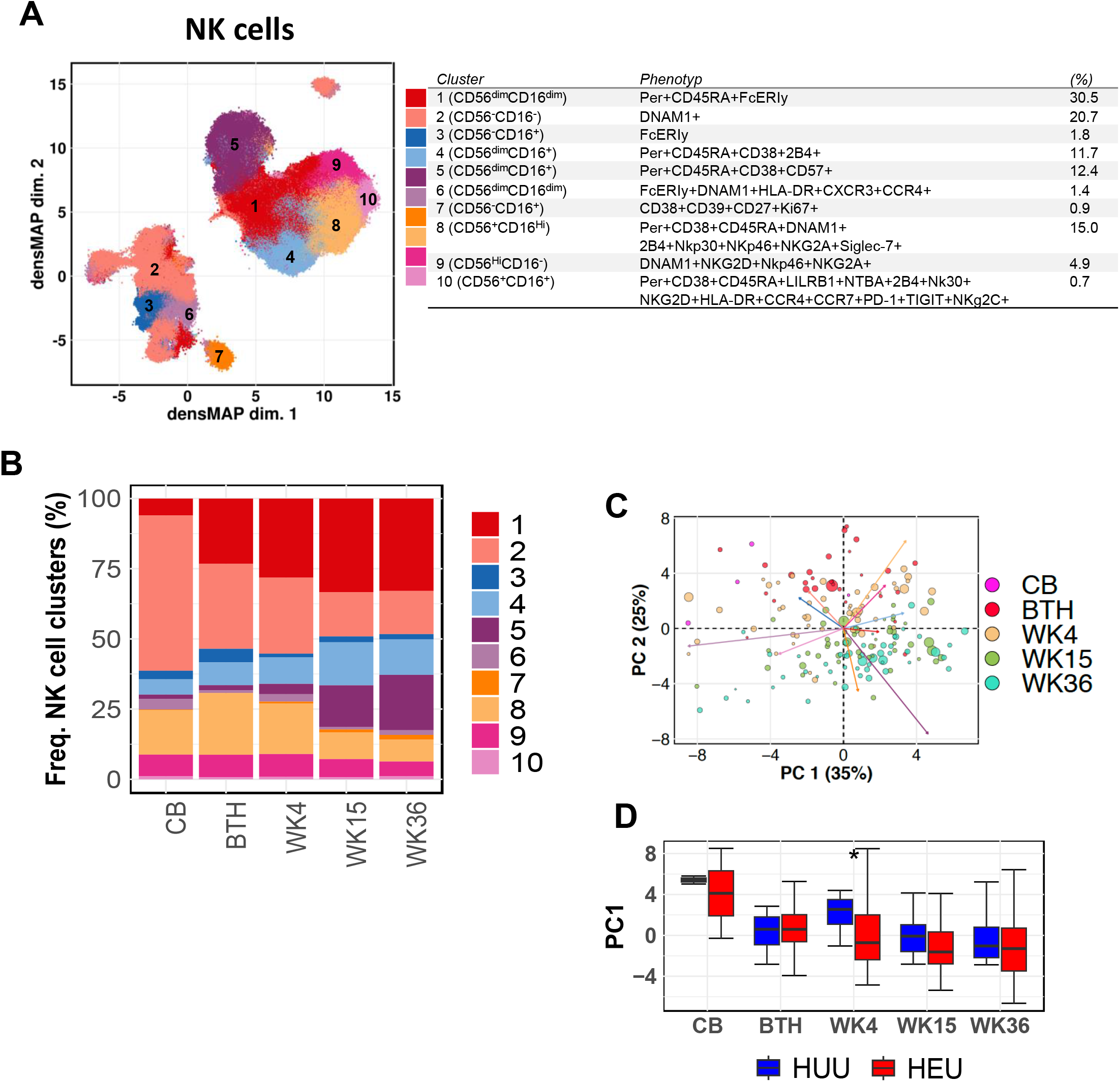
Early life immunophenotypic alteration in NK cells in HIV-exposed uninfected infants (iHEU) compared to HIV-unexposed uninfected infants (iHUU). **A)** Uniform manifold approximation and proximation (UMAP) with density preservation showing dimensional reduction of FlowSOM clusters on NK cells and table summarising immune cell cluster phenotype. **B)** Relative abundances of NK cell clusters measured in cord blood (CB) and infant peripheral blood at birth (BTH), weeks (WK) 4, 15 and 36. **C)** Age related phenotypic composition of NK cells depicted as principal component (PC) coordinates of centred log-odd ratios of FlowSOM clusters for each infant samples and arrows indicating contribution of each cell cluster in scatter of PC components. **D)** Boxplots comparing PC coordinates of NK cells between iHEU and iHUU

**Table 3:**
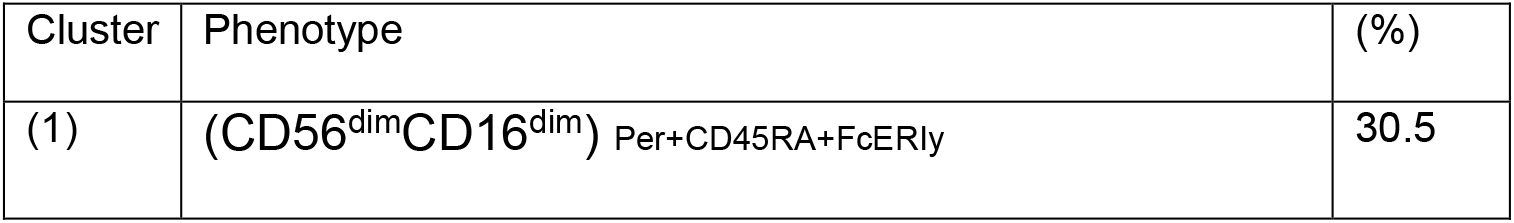
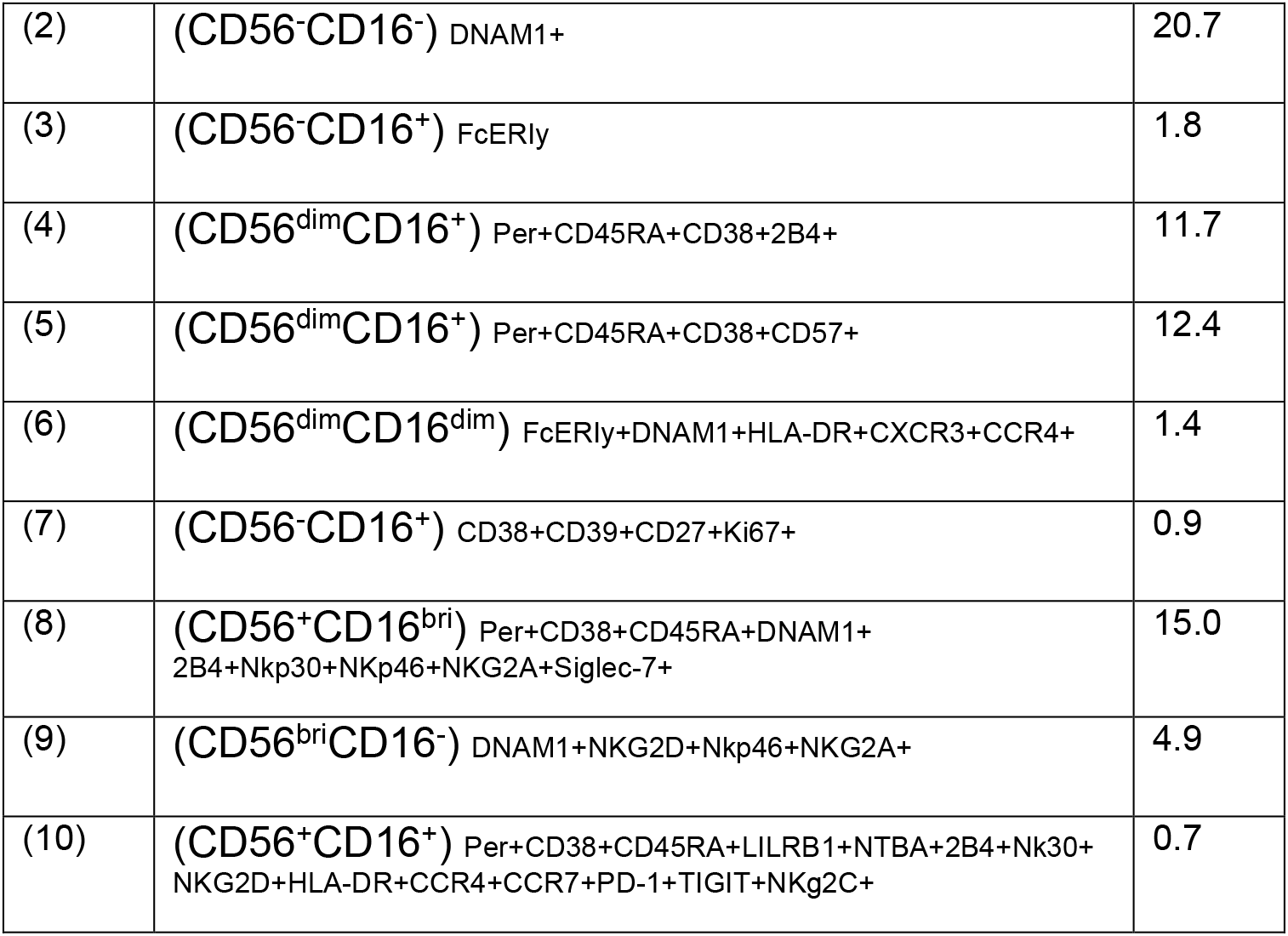
Phenotypic description of NK cell clusters identified by unsupervised cell clustering.

| Cluster | Phenotype | (%) |
| --- | --- | --- |
| (1) | (CD56<sup>dim</sup>CD16<sup>dim</sup>) Per+CD45RA+FcERly | 30.5 |
| (2) | (CD56<sup>-</sup>CD16<sup>-</sup>) DNAM1+ | 20.7 |
| (3) | (CD56<sup>-</sup>CD16<sup>+</sup>) FcERly | 1.8 |
| (4) | (CD56<sup>dim</sup>CD16<sup>+</sup>) Per+CD45RA+CD38+2B4+ | 11.7 |
| (5) | (CD56<sup>dim</sup>CD16<sup>+</sup>) Per+CD45RA+CD38+CD57+ | 12.4 |
| (6) | (CD56<sup>dim</sup>CD16<sup>dim</sup>) FcERly+DNAM1+HLA-DR+CXCR3+CCR4+ | 1.4 |
| (7) | (CD56<sup>-</sup>CD16<sup>+</sup>) CD38+CD39+CD27+Ki67+ | 0.9 |
| (8) | (CD56<sup>+</sup>CD16<sup>bri</sup>) Per+CD38+CD45RA+DNAM1+<br>2B4+Nkp30+Nkp46+NKG2A+Siglec-7+ | 15.0 |
| (9) | (CD56<sup>bri</sup>CD16<sup>-</sup>) DNAM1+NKG2D+Nkp46+NKG2A+ | 4.9 |
| (10) | (CD56<sup>+</sup>CD16<sup>+</sup>) Per+CD38+CD45RA+LILRB1+NTBA+2B4+Nk30+<br>NKG2D+HLA-DR+CCR4+CCR7+PD-1+TIGIT+NKG2C+ | 0.7 |

### Early TCR repertoire skewing in iHEU

To relate altered T cell memory maturation trajectories to T cell repertoire changes, we characterised TCR clonotypes in fractionated naïve and memory T cells from matching samples used to define memory lineage in the CyTOF panel. Each PBMC sample was sorted into four T cell subsets: naïve (CD45RA^+^CD27^+^CCR7^+^) CD4^+^ and CD8^+^ cells and total memory (CD45RA^-/+^CCR7^-/+^) T cells (Figure S3A). We subjected these sorted cell fractions (n=934 samples) to bulk sequencing of the TCRβ locus, which enabled identification of TCR clones in 885 samples. The number of unique clones identified per T cell subset positively correlated with the total number of cells sequenced (Figure S3B). There were 24 samples that had either fewer than 10 clones or the number of unique clones were greater than the number of cells sequenced, and these were removed from downstream analysis. Therefore, we included 861 samples having a total of 238,092 reads (range: 11-5451) and an overall median number of unique clones of 81 (IQR: 42-156). The CDR3 lengths of the clonotypes were evenly distributed across infant age and between iHEU and iHUU (Figure S3C).

The diversity and richness of the TCR clonotypes, determined by inverse Simpson diversity index and Chao1 richness scores respectively, remained unchanged with infant age (Figure S3D & S3E). However, when the TCR diversity scores were stratified by infant HIV exposure status, iHEU TCR clonotypes had relatively lower diversity compared to iHUU starting as early as birth (Figure 4A). These differences were apparent for naïve CD4^+^ T cells at weeks 15 and 36 and for naïve CD8^+^ T cells at birth and week 36. Differences in memory T cell diversity scores were observed to occur only at birth, being significantly lower in iHEU for both CD4^+^ and CD8^+^ T cells (Figure 4A). Similarly, richness (the number of unique TCR clones) was significantly lower in iHEU relative to iHUU for both CD4^+^ and CD8^+^ T cells at birth (Figure 4B). This remained statistically significant for naïve CD4^+^ T cells at week 36 (Figure 4B). When examining memory T cells, significant differences in TCR richness were restricted to earlier timepoints, being lower in iHEU compared to iHUU at birth and week 4 for CD4^+^ T cells and only at birth for CD8^+^ T cells. We also observed differences in the overall structural overlap, as measured by Jaccard indices, of the memory CD4^+^ TCR repertoire between iHEU and iHUU at birth, although this difference was less prominent at later time points (Figure S3F). In contrast, the TCR structural overlap for naïve CD8^+^ T cells was lower in iHEU compared to iHUU only at week 36 (Figure S3F). Differences in Vβ gene usage were observed between iHEU and iHUU with most genes being used more frequently in iHEU (Figure S4). Together, these results show expansion of specific TCR clones among iHEU, resulting in skewing of the TCR repertoire relative to iHUU.

**Figure 4:**
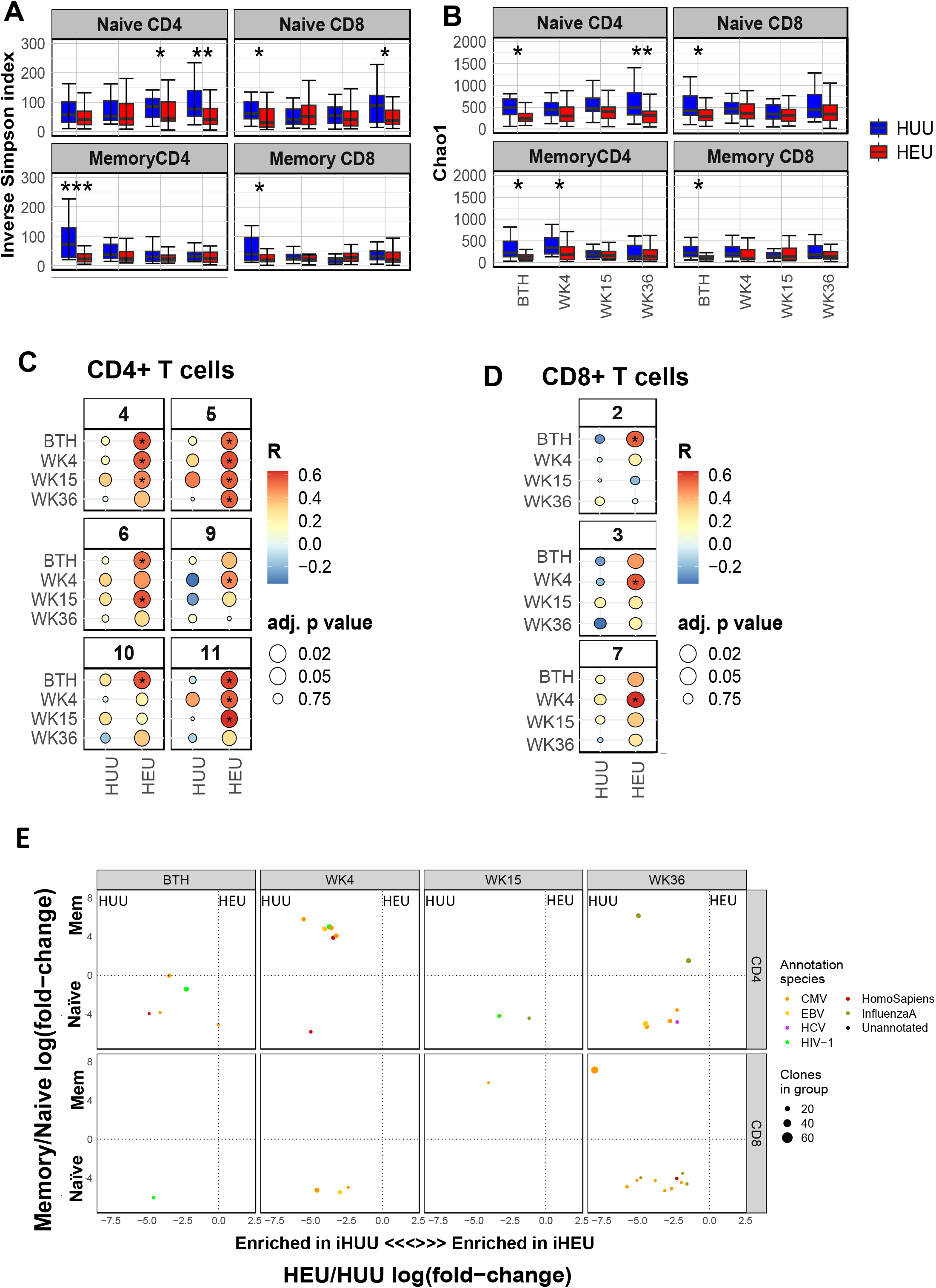
Premature CD4+ and CD8+ T cell receptor (TCR) repertoire skewing in HIV-exposed uninfected infants (iHEU) relative to HIV-unexposed uninfected infants (iHUU). **A)** Boxplots comparing Inverse Simpson TCR diversity scores between iHUU and iHEU at birth and weeks 4, 15 and 36. **B)** Boxplots comparing Chao1 TCR clonotype richness between iHUU and iHEU at birth and weeks 4, 15 and 36. **C &D**) Spearman’s rank correlation between naive CD4+ and CD8+ T cell Inverse Simpson scores and frequencies of FlowSOM clusters for CD4+ and CD8+ T cell clusters respectively. **E**). GLIPH analysis showing antigen specificity groups of the top 100 TCR clones that are significantly different between iHEU and iHUU and between naïve and memory CD4+ and CD8+ T cell subsets.

We next wished to determine if there was a relationship between TCR diversity and the clusters of CD4^+^ and CD8^+^ T cells identified in Figures 2A-F. By using Spearman’s rank correlation between T cell clusters and the inverse Simpson TCR diversity scores for naïve and memory CD4^+^ and CD8^+^ T cells, we found that the frequencies of CD4^+^ T cell clusters (4-6: CD38+HLA-DR^+^CD45RA^+^CD27^+^CCR7^+^, CCR4^+^CCR6^+^CXCR3^+^CD57^-^, CCR4^+^CCR6^+^CXCR3^+^CD57^+^, and 9-11: Perforin^+^Ki67^+^CD25^+^, Perforin^+^Ki67^-^HLA^-^DR^+^CD69^+^CD39^+^, Per^+^Ki67^+^) from iHEU were positively correlated with the TCR diversity scores derived from sorted naïve CD4^+^ T cells (Figure 4C). Similarly, frequencies of CD8^+^ T cells clusters (2, 3 and 7: CD56^+^CD16^+^NKp46+NKG2A^+^, HLA-DR^+^, Ki67^+^CD45RA^+^CD27^+^CCR7^-^) were positively correlated to the TCR diversity scores derived from the sorted naïve CD8^+^ T cells at either birth or week 4 (Figure 4D). No T cell clusters were significantly correlated to TCR diversity derived from sorted memory T cells (data not shown). Collectively, our findings show skewing of the TCR repertoire in iHEU relative to iHUU from birth, in the naïve T cell compartment prior to phenotypically defined memory maturation of CD4 and CD8 T cells.

### Predicted TCR recognition of epitopes found only in iHUU

To understand the predicted antigen specificities of the TCR clonotype differences between iHEU and iHUU, we used GLIPH (grouping of lymphocyte interactions by paratope hotspots)^27, 28^. TCR specificities from sorted naïve CD4^+^ T cells in iHUU appeared enriched with specificities for cytomegalovirus (CMV), Hepatitis C virus (HCV), SARS-Cov2 and HIV-1 (Figure 4E). Enrichment of Influenzae A specific clones was more apparent in memory CD4^+^ T cells at the week 36. For memory CD4^+^ T cells at week 4, the shared specificity groups in iHUU were enriched for CMV clones. Similarly, for CD8^+^ T cells, there were more specificities in the naïve compartment at both earlier and later time points with less predictions in memory CD8^+^ T cells. TCR specificities in sorted naïve CD4^+^ and CD8^+^ cells from iHUU appeared consistent throughout 36 weeks, whereas specificities in the CD4 memory compartment fluctuated and appeared predominantly at 4 and at 36 weeks (Figure 4E). Conversely for iHEU sorted T cell populations, no enriched TCR specificities in either naïve or memory CD4^+^ or CD8^+^ cells at any of the time points were predicted.

### NK cell clusters are the strongest predictors of antibody responses

To relate how early life T and NK cell clusters may associate with vaccine-induced antibody responses, we measured the response to acellular pertussis vaccination from birth to 36 weeks and to rotavirus at 36 weeks. We initially hypothesised that iHEU would have lower IgG and IgA responses compared with iHUU after vaccination with routine administration of these vaccines, which may explain a mechanism for higher infection rates of these pathogens in iHEU. To test this, IgG antibody levels against pertussis were measured at birth, weeks 4, 15 and 36, with rotavirus specific IgA and neutralization titres measured at week 36 (Figure 5B and C). The median anti-pertussis IgG OD values were lower in iHEU compared to iHUU at birth and week 4 (pre-vaccination at 6 weeks; Figure 5A). However, at week 15 (1 week post 3^rd^ vaccination), iHEU showed a significantly higher IgG response relative to iHUU, persisting until week 36 (Figure 5A). No significant differences were observed in anti-rotaviral IgA levels or ability to neutralize rotavirus between iHEU and iHUU (Figure 5B, C).

**Figure 5.**
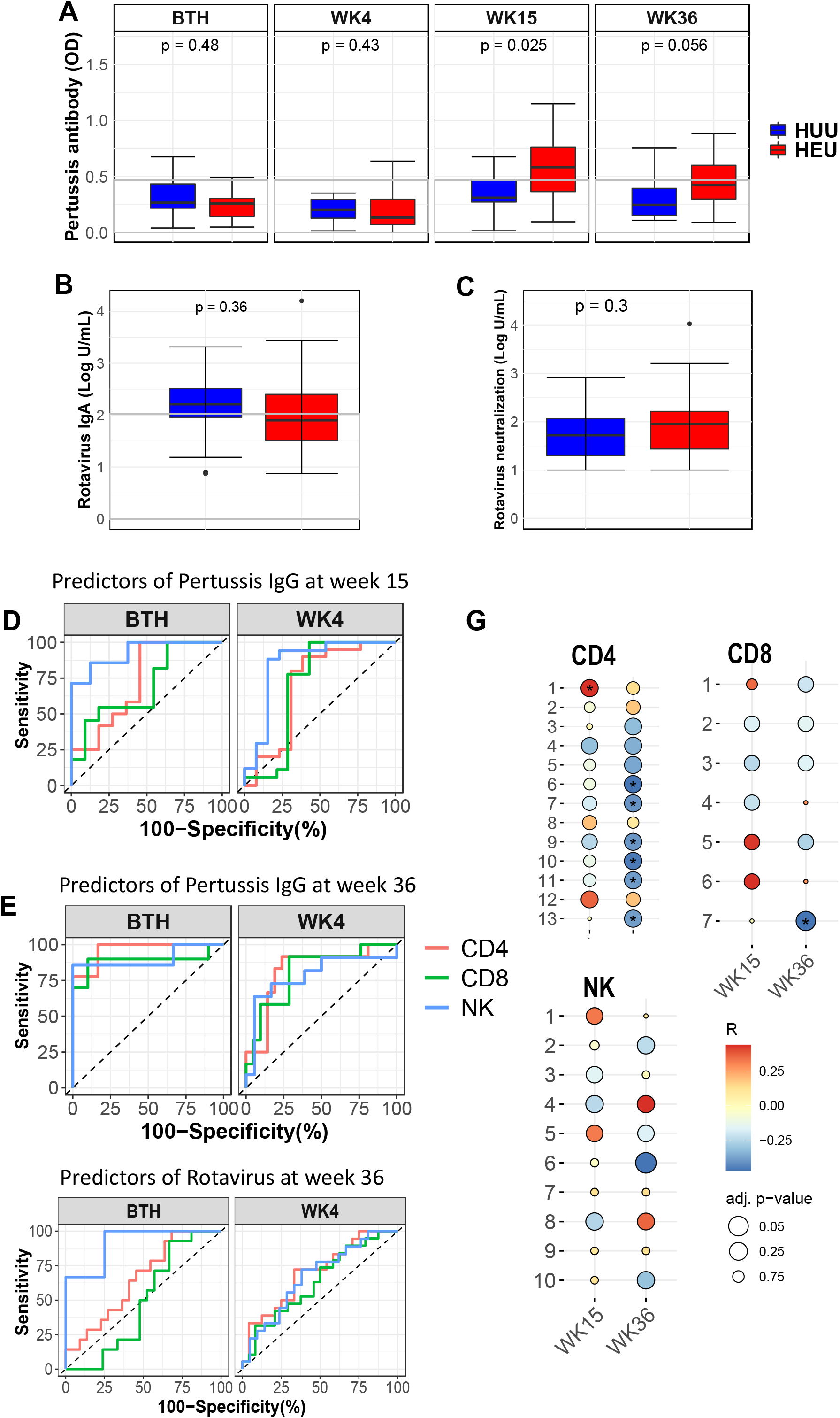
Association of vaccine antibody responses to immune cell phenotypes. **A)** Comparing IgG levels against pertussis between HIV-exposed uninfected infants (iHEU) and HIV-unexposed uninfected infants (iHUU), grey shaded area indicate threshold IgG levels for protective pertussis vaccine response. **B & C**) Boxplots comparing rotavirus specific IgA and neutralization titres between iHEU and iHUU at week 36. **D, E & F)** Summary of ROC analysis using the latent variable axis-1 derived from partial least square discriminate analysis (PLS-DA) of NK, CD4 and CD8 T cell clusters determined to be best predictors at birth and week 4 of pertussis antibody responses at weeks 15 and 36 and rotavirus antibody response at week 37. **G)** Spearman’s correlation between abundances FlowSOM clusters of CD4+ and CD8+ T cells and NK cells and anti-pertussis IgG titres measured at week 15 and 36.

We built a prediction model by grouping all infants (iHEU and iHUU) into those that had a protective pertussis-specific IgG titre (≥0.469 OD_Abs_) denoted as responders and those that were non-responders (<0.469 OD_Abs_). Most of the infants at birth and week 4 (89.7% and 94.1%, respectively) were determined to be non-responders as expected, since this was prior to vaccination. At week 15 and 36 of age, 46.4% and 34.0% infants respectively were determined to be responders (Figure 5A). Multivariate unbiased variable selection using MUVR^29^ was performed to determine early life cell clusters that could distinguish later pertussis vaccine responders from non-responders, according to our classification. Certain cell cluster abundances at birth and week 4 selected using MUVR were associated with low misclassification scores of a pertussis vaccine response at weeks 15 and 36 (Figure S4). We then used these abundances of the MUVR selected clusters to perform a partial least squares discriminant analysis (PLS-DA) with the pertussis IgG responses at week 15 and 36 as response variables (Figures 5D and E, respectively). The latent variables (LV) of the cell clusters as determined by PLS-DA were then used to compute area under the curve (AUC) of the receiver operating curve (ROC) distinguishing pertussis responders from non-responders. This approach revealed that NK cell clusters (1, 3, 4, 6, 8 and 10) at birth and at week 4 (clusters 1, 2, 3 and 9, Table 3) were ranked the highest in the list of predictors of the pertussis IgG vaccine response at week 15 (Figure 5D, S4A & S4B, at birth: AUC = 0.93 and p=0.005; at week 4: AUC = 0.85 and p=0.012). The cell clusters derived from CD4^+^ or CD8^+^ T cells were not significantly predictive of an IgG pertussis vaccine response at weeks 15 and 36 (Figure 5D, S4B-D, for CD4 clusters at birth: AUC=0.73, p=0.063; for CD8 clusters: AUC=0.69, p=0.12; at week 4: AUC=0.70, p=0.06; for CD8 clusters: AUC=0.70, p=0.053).

Using the same approach for rotavirus responses, we used an arbitrary threshold of overall median concentrations for IgA responses to dichotomize infants into those that had concentrations below the median (low-responders) and those above the median (high-responders; Figure 5B). In iHUU, 66.7% were high rotavirus vaccine responders compared to 45.5% iHEU (p=0.11). We subsequently determined which immune cell clusters at birth and week 4 were predictive of IgA responses measured at week 36 using MUVR (Figure 5F). PLS-DA analysis using the MUVR selected cell clusters revealed that NK cells at birth were highly predictive of an IgA rotavirus response at week 36 (Figure 5F, AUC = 0.92, p = 0.0007). Neither CD4^+^ or CD8^+^ T cell clusters were predictive of a rotavirus response at week 36 (Figure 5F, for CD4 clusters at birth: AUC=0.7, p=0.05; for CD8 clusters: AUC=0.55, p=0.59; at week 4: AUC=0.68, p=0.042; for CD8 clusters: AUC=0.6, p=0.24).

We also tested whether contemporaneous cell clusters might be related to pertussis responses. When we analysed the frequencies of the cell clusters at weeks 15 and 36 with pertussis antibody levels at weeks 15 and 36, CD4^+^ T cell cluster 1 (Th2 cells) was significantly correlated to pertussis antibody titres at week 15, while at week 36 CD4^+^ T cell clusters 6, 7, 9-11, and 13 (Table 1) were inversely correlated to pertussis antibody titres (Figure 5G). Only CD8^+^ T cell cluster 7 (Proliferating EM: Ki67^+^CD45RA^+^CD27^+^CCR7^-^) was observed to be inversely correlated to pertussis antibody titres measured at week 36 (Figure 5G). Of note, there was no significant contemporaneous correlation between the NK cell clusters and pertussis antibody titres (Figure 5G).

Collectively, these data show that various NK cell phenotypes at birth and in the first 1 month of life can predict vaccine-induced antibody responses to acellular pertussis and rotavirus at 9 months, regardless of infants being exposed to HIV.

## Discussion

Herein, we describe immune maturation in the first 9 months of life and how HIV/ARV exposure alters this ontological trajectory. Our findings show ordered immune changes during infancy, typical of the transition from innate-like and naïve neonatal immunity towards memory differentiated adaptive infant immunity^30^. NK cell phenotype and cluster composition differed between iHEU and iHUU at birth and week 4 and was predictive of vaccine responses later in life, whereas T cell memory maturation was impacted after 15 weeks. There was a skewing of the TCR repertoire in naïve and memory cells in iHEU from birth, relative to iHUU, and the persistent lack of TCR diversity that could account for the increased vulnerability to infections reported among iHEU^10^.

Although differentiated T cells are detected as early as 12-14 weeks in gestation^31, 32^, neonatal immunity is predominantly naïve and characterised by the abundance of CD45RA^+^ T cells^7^. T cell memory expansion during infancy is therefore essential in protecting against pathogens^33^. The differences in the kinetics of CD4^+^ and CD8^+^ T cell memory maturation in our study could not be explained by changes in TCR clonotypes since for both T cell subsets, TCR diversity and richness did not vary over time for either iHEU or iHUU. We presume that the steady rate of CD4^+^ T cell maturation is partially explained by the abundance of CD4^+^ T cells early in life, responsible for regulating immunological responses and thus facilitating adaptation to an extrauterine life laden with microorganisms^6, 34, 35^.

We show that the T cell compartment was significantly impacted by HIV/ARV exposure. The overall trajectory of T cell phenotypic composition in iHEU diverged from that of iHUU, particularly after week 15 of life. CD4^+^ and CD8^+^ T cell compartments in iHEU had lower frequencies of activated cells and a minor cell population expressing either perforin or PD-1 compared to iHUU. These findings contradict earlier reports showing iHEU having heightened immune activation and/or exhaustion^36, 37^, which may be related to mixedfeeding or non-breastmilk feeding that could introduce water contaminants or alter infant microbiome^38^. Mothers in our study were encouraged to exclusively breastfeed, likely accounting for lower levels of activated T cells in iHEU compared to iHUU^39^. Nonetheless, reduced or impaired memory T cells among iHEU compared to iHUU have been previously demonstrated and associated with poor childhood vaccine responses and/or increased hospitalization^16, 39–41^.

The 15 week period before significant expansion of T cell memory differentiation was preceded by significantly reduced diversity and richness in memory T cell clonality from birth in iHEU relative to iHUU. This was in agreement with others who reported lower βTCR clonality in the cord blood of iHEU^42^ and such reduced TCR diversity found at birth infers that T cells have undergone clonal expansion *in utero*. Using GLIPH^27, 28^, we could not detect any predicted antigen specificity enriched among iHEU, in contrast to earlier studies showing high frequency of HIV-1 specific clones^42^ and reactivity of T cells towards HIV proteins in iHEU^43^. What was evident instead was that naïve CD4^+^ and CD8^+^ TCR reactivities in iHUU were targeting CMV, EBV and HCV antigens. There may be three possible reasons for naïve cells showing clonal expansion and increased diversity in iHEU. Firstly, that our isolation of CD45RA+CCR7+CD27+ naïve population was contaminated with some memory cells during the sorting procedure. The contribution of sorting contamination would have negligible outcomes as we obtained >95% purity after sorting and any inclusion of memory cells would be of extremely low proportion. Additionally, the frequency of naïve T cells was significantly higher compared to that of memory T cells throughout infancy which could contribute to sampling bias resulting in enriched TCR clones amongst naïve T cells compared to the memory fraction. Second, that these cell populations are very early differentiated memory, and in the absence of CD95, we could not account for persistence of CD4 or CD8 memory stem cell emergence at birth^44–46^. The strong association between activated “naïve” CD4 and CD8 cells with TCR diversity would give credibility to this possibility. We hypothesised that such an outcome could be due either early differentiated cells that still expressed markers characteristic of naïve cells or the inclusion of stem cell-like memory cells that co-express CD27 and CD45RA^46^. Thirdly, these cells are not antigen-primed, but rather akin to virtual T cells^47, 48^. It has been shown that inflammatory conditions elicit virtual CD8^+^ T cells in the absence of specific antigen^49^. Further investigations are required to decipher the mechanism responsible for the diverse clonality found in naïve cells and whether inflammatory conditions in iHEU maybe driving this.

Unlike the somatically rearranged TCR repertoire, NK cell diversity arises through combinatorial expression of germline encoded receptors that sense expression of MHC class I molecules through surface bound KIR^23^. Previous viral encounters have been shown to augment NKR diversity and in some instances consequently reduce the cytolytic capacity of NK cells^50, 51^. We observed compositional differences of the NK cell clusters between iHEU and iHUU at week 4 in which iHEU had marginally higher memory-like/terminally differentiated NK cells (NKG2A^-^Per^+^CD57^+^) albeit not significantly so after adjusting for multiple comparisons. Enhanced cytolytic activity and elevated levels of the degranulation marker, CD107a, in iHEU compared to iHUU at birth have been previously reported^21^. Certain NK cell clusters at birth were superior in predicting both pertussis and rotavirus vaccine-induced IgG and IgA at 3-9 months compared to CD4+ T cells. Depending on the immunological milieu, NK cells can either negatively or positively regulate adaptive immunity, including promoting immunoglobulin isotype switching and enhancing antibody production^52, 53^. This finding was found for both iHEU and iHUU and mechanistic investigations would need to determine the role of NK cells in vaccine-induced antibodies.

The novelty of our study lies with the longitudinal follow-up of our infants with matched samples for multiple analyses allowing us to detail the ontogeny of T and NK cells from birth to 36 weeks of life and their relation with vaccination. The limitation of our study is the small sample volumes collected from the infants and our consequent inability to investigate functional changes in the immune subsets identified. A further limitation was the inability to tease out the effect of HIV from ARV exposure since all pregnant women living with HIV receive ARV treatment as a standard of care^54^.

In conclusion, our data show a transitional immunity during infancy, which is impacted by HIV/ARV exposure in a sequential manner starting with alteration of NK cells followed by T cell differentiation. Of particular importance, is the finding that TCR clonotypic diversity is significantly lower in iHEU from birth suggesting in utero skewing well before T cell memory formation. In addition, our data show that the composition of early life NK cells could predict vaccine responses in the first 9 months of life, although how this is accomplished will require further studies. Lastly, we show here a comprehensive phenotypic view of early life immune changes in response to HIV/ARV exposure. These changes may well be linked to the observed disparities in co-morbidities between iHEU and iHUU.

## Methods

### Study cohort

We analysed infants who were delivered by mothers living with and without HIV infection who were enrolled prospectively from birth and followed up until 9 months of age as previously described (Table S2)^55^. Briefly, pregnant women ≥18 years who recently delivered <12 hr were enrolled in the study together with their respective infants following signed informed consent. All mothers living with HIV received combined antiretroviral treatment during pregnancy under the Option B programme. The enrolment was restricted to infants having birth weight ≥ 2.5 kg, gestation ≥ 36 weeks and no complication experienced during delivery. Of those delivered by mothers living with HIV, viral transmission was assessed by performing HIV DNA PCR test after 6 weeks of life, and those who tested positive for perinatal HIV infection were excluded from further analysis. The infants participating in this study received childhood vaccines according to the South African Extended Program of Immunization (EPI). This included administration of acellular pertussis vaccine at weeks 6, 10 and 14 and rotavirus vaccine at weeks 6 and 14. Blood samples were collected at birth, weeks 4, 15 and 36 for isolation of plasma and peripheral blood mononuclear cells (PBMC). Demographic characteristics of the mothers and their respective infants are included in the study analysis and summarised in Table S2.

#### Plasma and peripheral blood mononuclear cell processing

Infant blood samples (0.5-3 mL) were collected into sodium heparin tubes and processed within 6hr. Plasma and PBMC were isolated using ficoll centrifugation. Plasma samples were stored at -80°C while PBMC were cryopreserved in 90% Fetal Calf serum (FCS) with 10% DMSO in liquid nitrogen.

#### Intracellular and surface staining of PBMC for mass cytometry

A total of 278 infant samples (including 27 duplicates) were used to assess the longitudinal changes of immune cells using mass cytometry and TCR sequencing. PBMC samples were retrieved from liquid nitrogen and thawed at 37°C before being transferred into RPMI 1640 media supplemented with 10% FCS and 10KU Benzonase. Cells were centrifuged, washed twice in PBS and counted using a TC20 cell counter (Biorad). We aimed to stain 2x10^6^ viable cells for mass cytometry, however cell recovery varied by infant and age, with fewer cells collected at earlier time points due to smaller blood volumes. PBMC samples were split into 2 aliquots with 2x10^6^ (or ¾ for cells with <2x10^6^) used for mass cytometry staining and 1x10^6^ (or ¼ for cells with <2x10^6^) used for TCR sequencing. Lyophilised surface and intracellular antibody mixtures^56^ stored at 4°C were reconstituted in CyFACS buffer (PBS, 2% FCS) and permeabilization buffer (eBioscience permwash) respectively for cell labelling. Table S1 lists the antigen targets included in the mass cytometry antibody panel. Prior to antibody labelling, the cells were stained with cisplatin in PBS to determine cell viability followed by staining with surface antibodies for 20 min at room temperature. Cells were then fixed with 2% paraformaldehyde solution, permeabilised and stained with intracellular antibodies at 4°C for 45 min. Upon which cells were washed with CyFACS buffer and resuspended with 2% paraformaldehyde solution containing Iridium DNA intercalator overnight at 4°C. Samples were then washed with PBS and resuspended in MilliQ water containing EQ Four Element calibration beads (10% v,v, Fluidigm) prior to acquisition using CyTOF 2 instrument (DVS Sciences).

#### Fluorescent activated cell sorting of naïve and memory T cells

The remaining PBMC from each sample were used to sort for naïve and memory CD4^+^ and CD8^+^ T cells using fluorescent activated cell sorting. Cells were labelled by surface staining using T cell markers including CD3, CD4 and CD8 and memory markers CD27, CD45RA and CCR7 (Table S2 & Figure S3). Naïve cells were denoted as those co-expressing CCR7, CD45RA and CD27 while the other remaining cells were regarded as memory cells. A BD FACSAria Fusion (BD) was used for 4-way sorting of CD4 and CD8 naïve and memory T cells from each sample and collected directly into FCS. Sorted cells were centrifuged, resuspended in cell RNAProtect (Qiagen) and stored at -40°C until processing for TCR sequencing.

### Bulk TCR sequencing

RNA was purified from each sorted sample using the RNAeasy Plus Micro kit (Qiagen) and libraries were prepared for TCR sequencing as described by Rubelt, et al.^57^. This was performed at the University of Cape Town. Briefly, purified RNA was reverse transcribed in a Rapid amplification of cDNA ends (RACE) reaction using SMARTScribe Reverse Transcriptase using previously designed oligos (isoC-5′-GTCAGATGTGTATAAGAGACAGnnnnnnnnnnCGATAGrGrGrG -3′-C3_Spacer and for the Poly A tail 5′-GTGTCACGTACAGAGTCATCtttttttttttttttttttttttttttttt -3′ VN) that captures polyadenylated transcripts and introduces a 10bp unique molecular identifier (UMI) at the 5’ end of the product. cDNA was purified with AmpureXP beads (Beckman Coulter) before whole transcriptome amplification using Advantage 2 Polymerase (Clontech) using oligos that introduce Illumina Nextera Multiplex Identifier (MID) P5 Adapter sequences. The libraries were then sent to Stanford University (Department of Microbiology and Immunology, Stanford University) in a blinded fashion, where TCRβ-specific amplification was achieved with Q5 Hot Start Master Mix (NEB) using constant region-specific oligos, simultaneously introducing P7 MID sequences. Final sequencing libraries were purified using SPRISelect beads (Beckman Coulter), quantified using the Agilent TapeStation and pooled for sequencing. Paired-end sequencing was performed on an Illumina NovaSeq SP with 2x250 cycles, performed by the Chan-Zuckerberg Biohub Initiative.

### Quantification of vaccine antibody responses

Infant pertussis specific antibody responses were measured at birth, 4, 15 and 36 weeks of age using a commercial human IgG ELISA kit (Abcam). Plasma samples were diluted 1:100 into sample diluent antibody titres measured in duplicates according to manufacture instructions.

Rotavirus IgA titres were determined by enzyme immunoassay (EIA) and performed in a blinded manner in the Division of Infectious Disease, Department of Pediatrics, Cincinnati Childrens’ Hospital Medical Centre, Cincinnati, Ohio. Briefly, purified rabbit anti-rotavirus IgG were immobilised on microtiter plate as capture antibodies. Lysates from rotavirus (strains RV3 and 8912) and mock infected cells (control) were added to the immobilised capture antibodies to bind rotavirus antigens and any uncaptured antigens were washed off with PBS, 0.05% Tween20. Reference standards, control and test samples were diluted in PBS, 0.05% Tween20, 1% non-fat dry milk and 50 uL added to microtiter plate for antigen binding. Bound antibodies were detected by biotyinylated goat anti-human IgA (Jackson Laboratories) and the addition of peroxidase conjugated avidin:biotin (Vector Laboratories, Inc., Burlingame, CA). Chromogenic signal was generated by addition of substrate O-phenylenediamine (Sigma) and reaction stopped after 30 min using 1M H_2_SO_4_. Absorbance was measured at 492_nm_ on a Molecular Devices SpectraMax Plus plate reader. The reference standard was assigned 1000 arbitrary units (AU) and a four-parameter logistic regression was used to extrapolate anti-rotavirus IgA tires using SoftMax software.

Rotavirus neutralisation titres were determined as previously described^58^. In this study we used Wa G1P8, 1076 G4P6 and DS-1 G2P4 virus strains obtained from the National Institute of Health (NIH). Plasma samples were serially diluted and incubated with the rotavirus strain for neutralization prior to adding the mixture to susceptible MA104 (monkey kidney) cell line for overnight incubation. Cell lysates were used to determine the level of un-neutralized rotavirus antigens using the EIA as described using guinea pig anti-rotavirus antiserum to measure captured rotavirus antigens. Rabbit anti-guinea pig IgG conjugated to horseradish peroxidase (Jackson ImmunoResearch) was used to detect bound antibodies, and chromogenic signal generated using OPD. The amount of rotavirus present in the resuspended lysate from each well was inversely related to the amount of neutralizing antibody present in the plasma. Each plasma dilution series as modelled using a logistic regression function. For each fitted curve the dilution which corresponds to a 40% response (ED_40_), compared to the virus controls, was determined and reported as the neutralization titer. The ED_40_ represents the titre of the serum against a given virus, which represents a 60% reduction in amount of virus.

#### Data analysis

### Mass cytometry

#### Processing

Post CyTOF acquisition, the files were normalized using Premessa R package and target populations were manually gated using FlowJo software (version 10.5.3, TreeStar). The target populations were exported as FCS files for downstream analysis using R, an open-source statistical software^59^. All files for the target populations were transformed by applying inverse hyperbolic sine with a cofactor of 5 and subjected to doublet detection and removal using computeDoubletDensity from the scDblFinder package^60^.

#### Dimensional reduction

Median marker expressions for each sample were used to compute Euclidean distance matrix to determine multidimensional scaling (MDS) coordinates using the cmdscale function in R. MDS analysis were used to visualize immune cell trajectories. Further, marker expression intensities were used to implement Uniform Manifold Approximation and Projection (UMAP) while preserving local density distribution for visualization using densMAP function from the denvis R package^61^.

#### Cell Clustering

High resolution clustering of the target cell populations was performed using FlowSOM algorithm and the resulting clusters grouped into metaclusters using ConsensusClusterPlus available in the CATALYST R package^62^. The immune cell clusters were visualised using hierarchical clustering and UMAP embedding. Centred log-ratios of the relative abundance of the cell clusters per sample were determined and used in Principal Component Analysis (PCA) for assessing immune cell compositional differences.

## TCR sequencing

### Pre-processing of TCR sequencing data

Raw sequencing data was pre-processed by pRESTO^63^. Briefly, reads with mean Phred quality scores less than 20 were discarded, UMIs were extracted from the first 10bp of read 1, and primer sequences masked. Next, a single consensus sequence was constructed from reads sharing the same UMI barcode and paired-end consensus sequences assembled into a full-length TCR sequence. In the case of non-overlapping mate-pairs, the human TRB reference as used to properly space non-overlapping reads. After removal of duplicate sequences, full-length reads were aligned and clonotypes assembled using MiXCR^64, 65^.

### T cell immune repertoire analysis

To perform quality control on our bulk TCR sequencing data, we removed samples where fewer than 10 clones were detected (*n =* 44 samples) and where more clones were detected than cells were sorted (*n* = 18 samples). This yielded a total of 861 samples for downstream analysis. The immunarch packag^66^ operating in the open-source statistical software R was used for all downstream immune repertoire analysis, including calculation of CDR3β lengths and repertoire diversity by the Inverse Simpson index and richness using Choa1.

### Identification and annotation of T cell specificity groups

The GLIPH2 algorithm was used to establish T cell specificity groups, clusters of CDR3 sequences that are predicted to bind to the same antigen, by discovering groups of CDR3β sequences that share either global or local motifs^27, 28^. To assign specificity annotations to identified CDR3β sequence clusters, we adapted an approach described by Chiou, et al^67^. Briefly, we collected human CDR3β sequences with known specificity from VDJdb (https://vdjdb.cdr3.net/) and identified CDR3β sequences within this database that could form TCR specificity groups. From 47,107 CDR3β sequences, this process identified 11,648 specificity groups comprised of 25,759 unique CDR3β sequences. We next combined these 25,759 annotated sequences with 108,183 unique experimental CDR3β sequences from our bulk TCR repertoire profiling cohort. This was performed separately for CD4 and CD8 T, as there are differences between gene usage frequencies between CD4 and CD8 T cells that can impact specificity predictions. We prioritized TCR specificity groups with at least 3 distinct CDR3β sequences. This process yielded 17,154 CDR3β sequences in 11,629 specificity groups for CD4 T cells, and 15,107 CDR3β sequences in 9,911 specificity groups for CD8 T cells.

## Statistical analysis

All statistical analysis and graphical visualization of the data was performed on the open R software^59^. Spearman’s rank correlation was used to test for the correlations between the frequencies of the cell clusters with either infant age or antibody responses and p-values adjusted for multiple comparisons using false discovery rates (FDR). To compare differences in the abundance of cell clusters between iHEU and iHUU the generalised linear model from the diffcyt package was used^68^. Pairwise comparisons of medians between groups were performed using Wilcoxon-rank test or Kruskal-Wallis for multiple group comparisons and the p-values adjusted for multiple comparisons using Benjamin Hochberg correction. Generalised linear mixed model (GLMM) with bootstrap resampling from the R package CytoGLMM^69^ was used to identify NK cell markers that are predictive of HIV exposure at birth. To determine cell clusters that were predictive of antibody responses following vaccination, we used the multivariate modelling algorithm from MUVR that incorporates recursive variable selection using repeated double cross-validation within partial least squares modelling^29^.

## Acknowledgements

We would like to acknowledge the technical staff that assisted in the completion of this study. This includes Euan Johnston, Emily Tangie and Carine Kilola who assisted with the CyTOF experiments, Mikayla Stabile, Michelle Leong, Kassandra Pinedo, and Nicole Prins, for assistance with TCR RNA-seq processing, and Llyod Leach for assistance with measuring pertussis antibody responses.

## Author contribution

Conceived by CMG, CAB and HBJ, Method set-up and validation by SD, AW, SC, TR, FR, HH, and MD. Study participant enrolment, sample collection and processing by HBJ and BA. Experimental investigations by SD. Data and statistical analysis by SD, AW and SH. Original draft by SD. Reviewed and edited by all authors.

