## Supplementary material for "Disrupted memory T cell expansion in HIV-exposed uninfected infants is preceded by premature skewing of T cell receptor clonality": Suplementary information (Tables and Figures)

Supplementary information

Supplementary Tables

- Supplementary **Table S1** List of key resource materials used in the study.
- Supplementary **Table S2** Demographic characteristic of HIV-uninfected and HIV-infected mothers and their respective HIV uninfected-unexposed infants (iHUU) and HIV-exposed uninfected infants (iHEU).

Supplementary Figure

- **Figure S1** Immunophenotype of lineage cell clusters. **A)** Gating strategy used to determine live singlet cells of CD4+ and CD8+ T cells and NK cells subsets. **B)** Heatmap showing marker expression for FlowSOM immune lineage clusters derived from live singlet cells.
- **Figure S2** Immunophenotype of CD4+ and CD8+ T cell and NK cell FlowSOM clusters. **A)** Heatmap showin scaled marker expression for cell clusters derived from CD4+ and CD8+ T cells and NK cells. **B)** Generalized linear mixed model (GLMM) comparing relative abundance of CD4+ T cell clusters between HIV-exposed uninfected infants (iHEU) and HIV-unexposed uninfected infants (iHUU) at birth and week 4. **C)** GLMM comparing relative abundances of NK cell clusters between iHEU and iHUU at week 4 and 15.
- **Figure S3** Naïve and memory CD4+ and CD8+ T cell receptor (TCR) repertoire in HIV-exposed uninfected infants (iHEU) and HIV-unexposed uninfected infants (iHUU). **A)** Flowplots showing gating strategy for sorting naïve and memory CD4+ and CD8+ T cells in infants peripheral blood mononuclear cells prior TCR RNA sequencing. **B)** Heatmap showing correlation matrix of TCR quality control parameters. **C)** CDR3 lengths distribution between iHUU and iHEU. **D)** Longitudinal changes in TCR diversity scores measured by Inverse Simpson index. **E)** Longitudinal changes in TCR richness scores measured by Choa1 index. **F**) Comparing TCR repertoire structure measured using Jacard indeces between iHEU and iHUU.
- **Figure S4** Comparing TCR Vb gene usages between HIV-exuposed uninfected infants (iHEU) and HIV-unexposed uninfected infants in CD4+ and CD+8 T cells measured at birth (BTH) and weeks (WK) 4, 15 and 36.
- **Figure S5** Immune cell clusters predictive of pertussis specific antibody responses post-vaccination. Multivariable regression using partial least

squares with discriminate analysis (PLS-DA) and recursive variable elimination within repeated double cross-validation for selection of the minimum number of cell clusters with low misclassification error for predicting pertussis specific IgG responses at week 15 and 36. **A & B)** NK cell clusters predictors at birth and week 4 respectively. **C & D) C**D4+ T cell predictors at birth and week 4 respectively. **E & F) C**D8+ T cell predictors at birth and week 4 respectively.

- **Figure S6** Immune cell clusters predictive of rotavirus specific antibody responses post-vaccination. Multivariable regression using partial least squares with discriminate analysis (PLS-DA) and recursive variable elimination within repeated double cross-validation for selection of the minimum number of cell clusters with low misclassification error for predicting rotavirus specific IgG responses at week 36. **A & B)** NK cell clusters predictors at birth and week 4 respectively. **C & D) C**D4+ T cell predictors at birth and week 4 respectively. **E & F) C**D8+ T cell predictors at birth and week 4 respectively.

Table S1 List of key resource materials

| **Mass cytometry antibodies** | |
|---|---|
| Antigen | Metal isotope |
| *Extracellular antibodies* | |
| CD19-CD20 | In115Di |
| CD14 | Nd150Di |
| CD3 | Nd142Di |
| CD4 | Tb159Di |
| CD8 | Nd144Di |
| CD45RA | Nd148Di |
| KIR2DL1 | Sm149Di |
| CD57 | Eu151Di |
| Siglec-7 | Eu153Di |
| PD-1 | Sm154Di |
| NKp46 | Gd155Di |
| NKG2D | Gd156Di |
| NKG2C | Gd157Di |
| 2B4 | Gd158Di |
| CXCR3 | Gd160Di |
| NKp30 | Dy161Di |
| CD39 | Dy162Di |
| KIR3DL1 | Dy163Di |
| TIGIT | Dy164Di |
| CD16 | Ho165Di |
| CD69 | Er166Di |
| CD127 | Er167Di |
| CCR7 | Er168Di |
| NKG2A | Tm169Di |
| KIR2DL3 | Er170Di |
| CCR4 | Yb171Di |
| NTBA | Yb172Di |
| CCR6 | Yb173Di |
| CD56 | Yb174Di |
| CD25 | Lu175Di |
| CD38 | Yb176Di |
| CD7 | La139Di |
| DNAM1 | Pr141Di |
| LILRB1 | Nd143Di |
| CD27 | Nd146Di |
| HLA-DR | Cd112Di |
| *Intracellular antibodies* | |
| FcERIγ | Nd145Di |
| Ki67 | Sm152Di |
| Perforin | Sm147Di |
| *Live/Dead and DNA markers* | |
| Cisplatin | Pt |
| DNA intercalator | Ir |
| **Fluorescent Activated Cell Sorting** | |
| Antigen | Fluorochrome |
| CD3 | FITC |
| CD4 | Alexa Flour 700 |
| CD8 | BV711 |
| CD45RA | PE-Texas red |
| CD27 | PE-Cy5 |
| CCR7 | PE-Cy7 |

| T cell receptor RNA sequencing |
|---|
| RNAProtect solution |
| RNAeasy Plus Micro Kit |
| SMARTScribe Reverse Transcriptase |
| Q5 Hot Start Master Mix |
| Advantage 2 Polymerase |
| Primers |
| isoC-5′-GTCAGATGTGTATAAGAGACAGnnnnnnnnnnCGATAGrGrGrG -3′-C3_Spacer |
| Poly A tail 5′- GTGTCACGTACAGAGTCATCtttttttttttttttttttttttttttttt -3′ VN |
| **ELISA antibody quantification** |
| Anti-Pertussis IgG ELISA Kit |
| Anti-Rotavirus IgA rabbit antibodies |
| Rotavirus strains RV3 and 8912 |
| biotyinylated goat anti-human IgA |
| peroxidase conjugated avidin:biotin |
| O-phenylenediamine |
| **Reagents** |
| Histopague Ficoll |
| Fetal Calf Serum |
| Benzonase |
| EQ Four element calibration beads |
| eBioscience permeabilization buffer |
| Paraformaldehyde |

Table S2 Demographic characteristic of HIV-uninfected and HIV-infected mothers and their respective HIV uninfected-unexposed infants (iHUU) and HIV-exposed uninfected infants (iHEU).

| Demographics | iHUU (N=16) | iHEU (N=40) | P |
|---|---|---|---|
| **Mother** | | | |
| Median maternal age at delivery, years (range) | 23.5 (19-39) | 30.5 (19-39) | 0.02 |
| Median gestational age, weeks (range) | 39.0 (36-41) | 39.0 (36-42) | 0.9 |
| Median maternal CD4 count cells/μL (IQR) | | 418.0 (285.0-526.0) | |
| Median maternal viral load copies/mL (IQR) | | 20.0 (20.0-70.0) | |
| **Infant** | | | |
| Median birth weight, kg (IQR) | 3.2 (2.8-3.5) | 3.1 (2.9-3.4) | 0.8 |
| Gender, female (%) | 8 (50.0) | 14(35.0) | 0.4 |
| Median duration of breastfeeding (IQR) | 44.3 (36.0-52.0) | 52 | 0.1 |
| Median duration of exclusive breastfeeding, weeks (IQR) | 12.5 (9.3-27.0) | 17.5 (6.3-36.0) | 0.8 |
| Completed rotavirus vaccination (%) | 13 (81.3) | 35 (87.5) | 0.7 |
| Completed pertussis vaccination (%) | 12 (75.0) | 33 (82.5) | 0.7 |

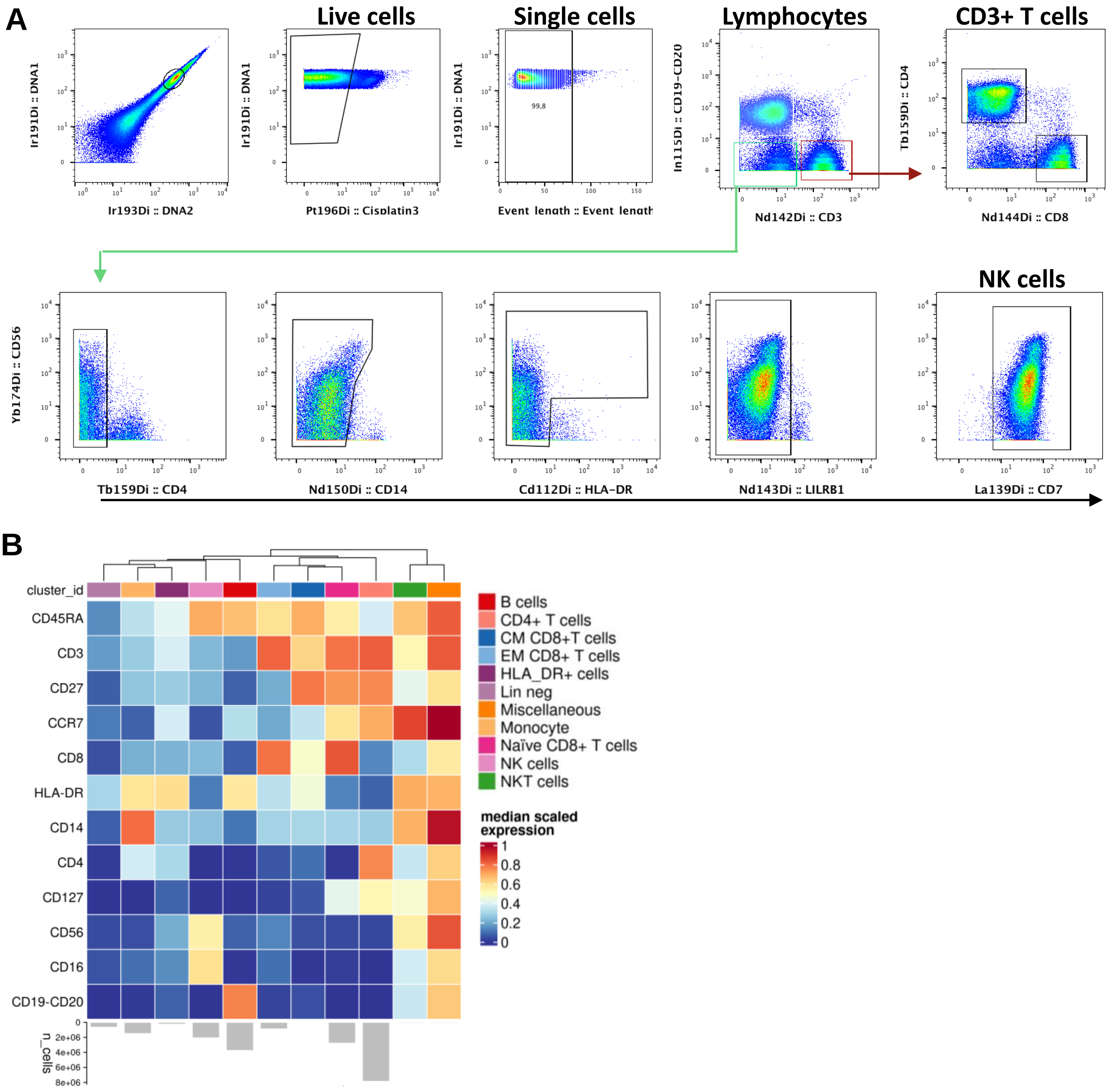

**Figure S1** Immunophenotype of lineage cell clusters. **A)** Gating strategy used to determine live singlet cells of CD4+ and CD8+ T cells and NK cells subsets. **B)** Heatmap showing re-scaled marker expression for FlowSOM immune lineage clusters derived from live singlet cells.

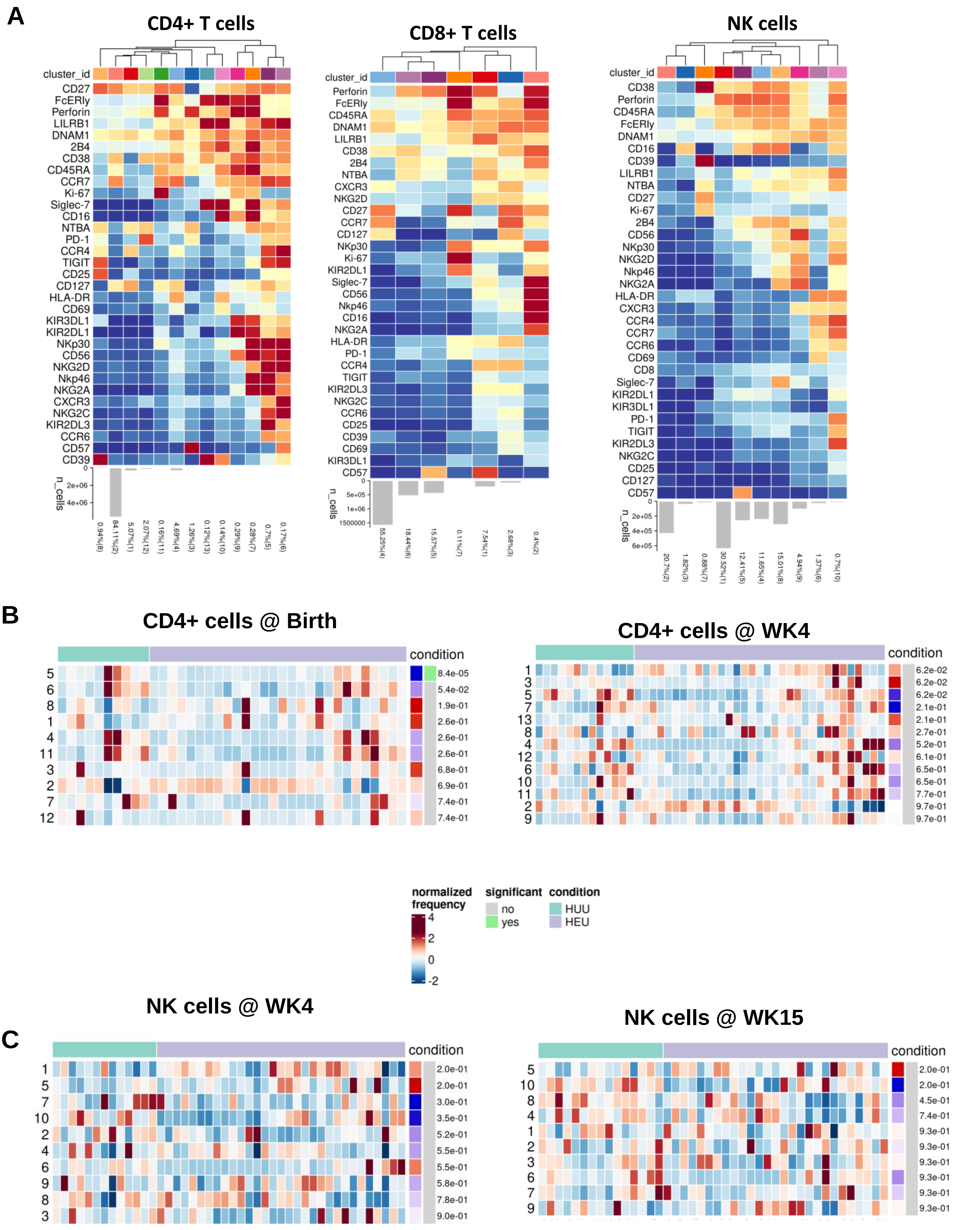

**Figure S2** Immunophenotype of CD4+ and CD8+ T cell and NK cell FlowSOM clusters. **A)** Heatmap showing re-scaled marker expression for cell clusters derived from CD4+ and CD8+ T cells and NK cells. **B)** Generalized linear mixed model (GLMM) comparing relative abundance of CD4+ T cell clusters between HIV-exposed uninfected infants (iHEU) and HIV-unexposed uninfected infants (iHUU) at birth and week 4. **C)** GLMM comparing relative abundances of NK cell clusters between iHEU and iHUU at week 4 and 15.

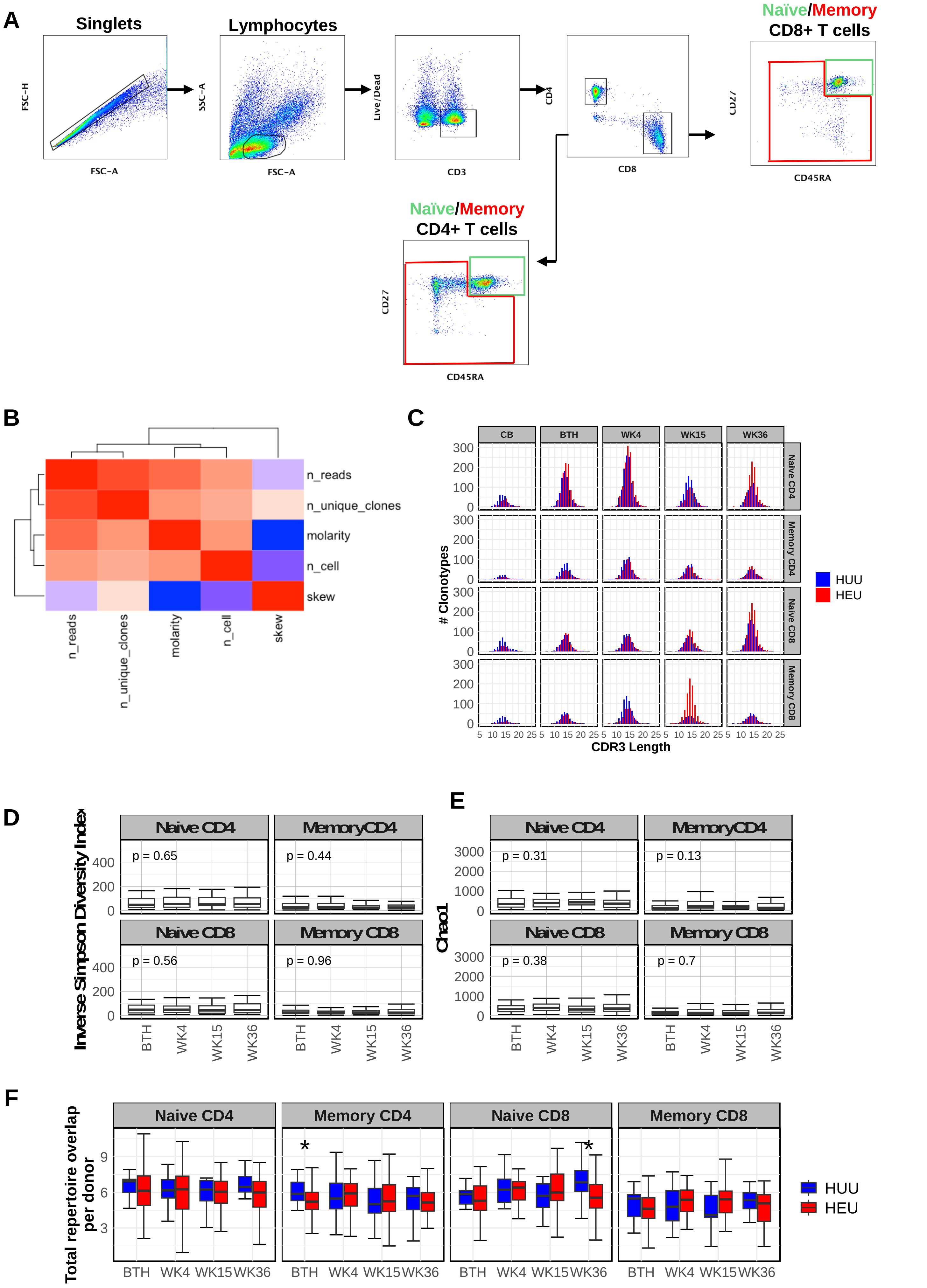

**Figure S3** Naïve and memory CD4+ and CD8+ T cell receptor (TCR) repertoire in HIV-exposed uninfected infants (iHEU) and HIV-unexposed uninfected infants (iHUU). **A)** Flowplots showing gating strategy for sorting naïve and memory CD4+ and CD8+ T cells in infants' peripheral blood mononuclear cells prior TCR RNA sequencing. **B)** Heatmap showing correlation matrix of TCR quality control parameters. **C)** CDR3 lengths distribution between iHUU and iHEU. **D)** Longitudinal changes in TCR diversity scores measured by Inverse Simpson index. **E)** Longitudinal changes in TCR richness scores measured by Choa1 index. **F)** Comparing TCR repertoire structure measured using Jacard indices between iHEU and iHUU.

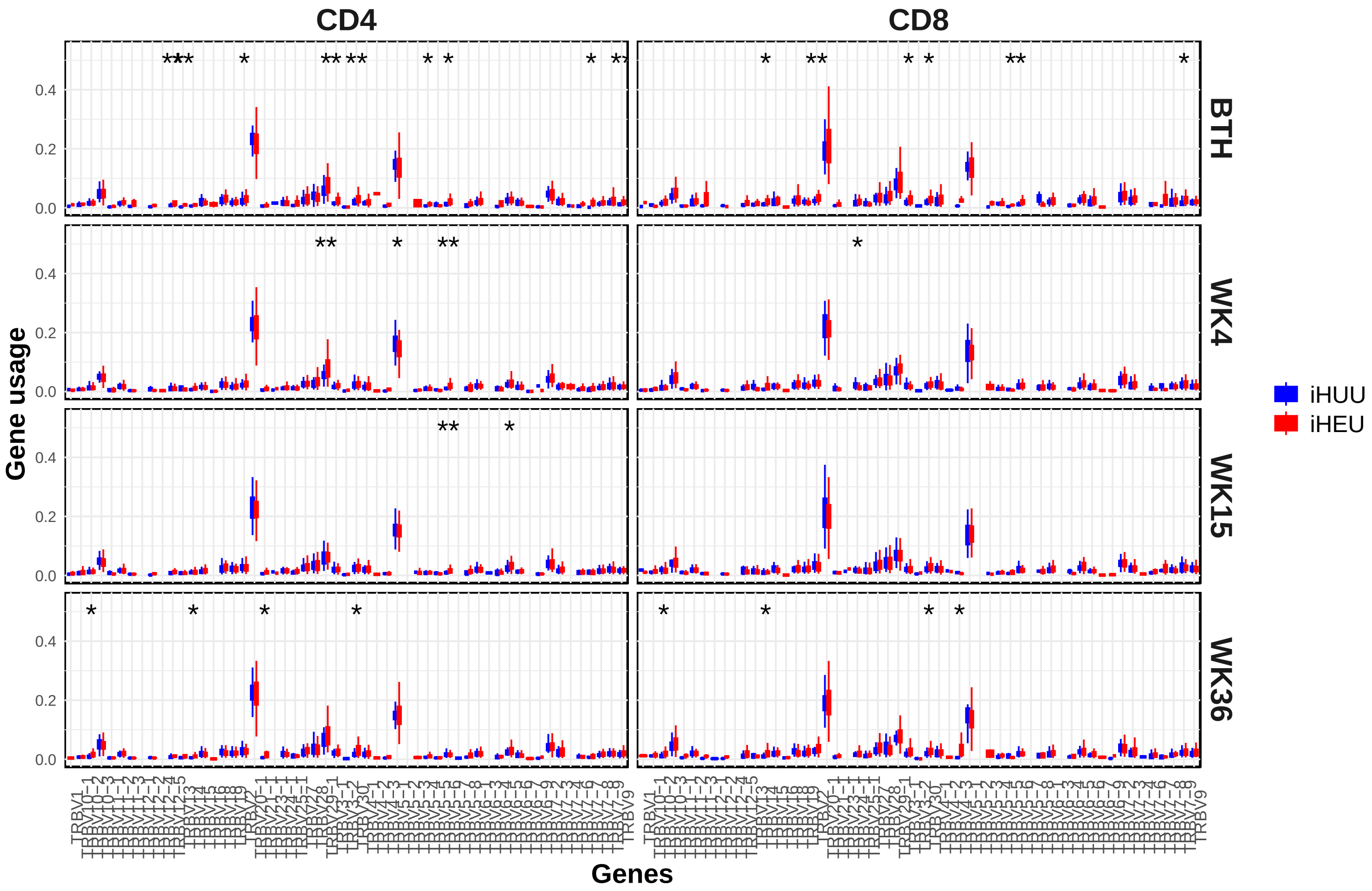

**Figure S4** Comparing TCR Vβ gene usages between HIV-exposed uninfected infants (iHEU) and HIV-unexposed uninfected infants in CD4+ and CD+8 T cells measured at birth (BTH) and weeks (WK) 4, 15 and 36.

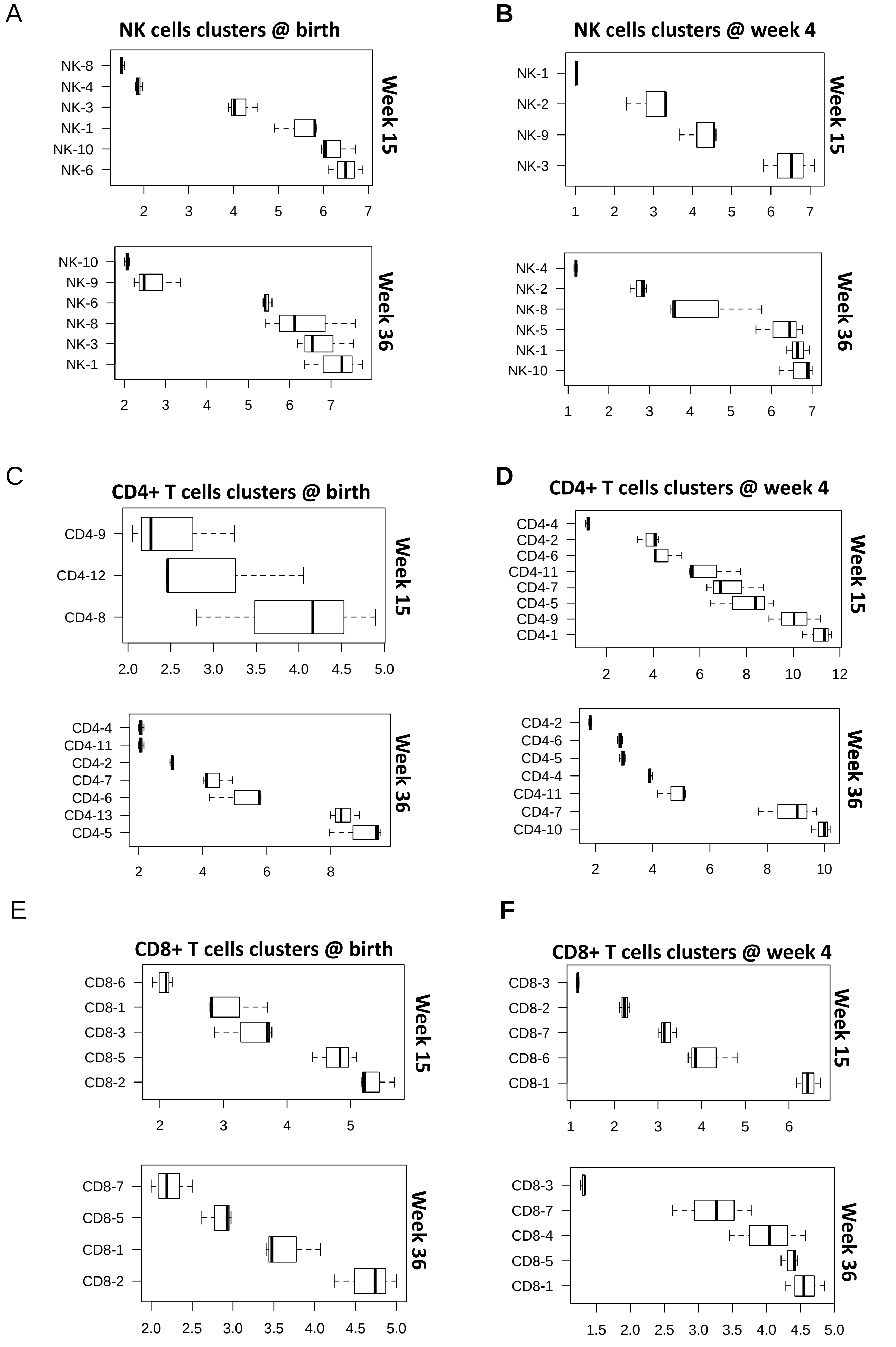

**Figure S5** Immune cell clusters predictive of pertussis specific antibody responses post-vaccination. Multivariable regression using partial least squares with discriminate analysis (PLS-DA) and recursive variable elimination within repeated double cross-validation for selection of the minimum number of cell clusters with low misclassification error for predicting pertussis specific IgG responses at week 15 and 36. **A & B)** NK cell clusters predictors at birth and week 4 respectively. **C & D) C**D4+ T cell predictors at birth and week 4 respectively. **E & F) C**D8+ T cell predictors at birth and week 4 respectively.

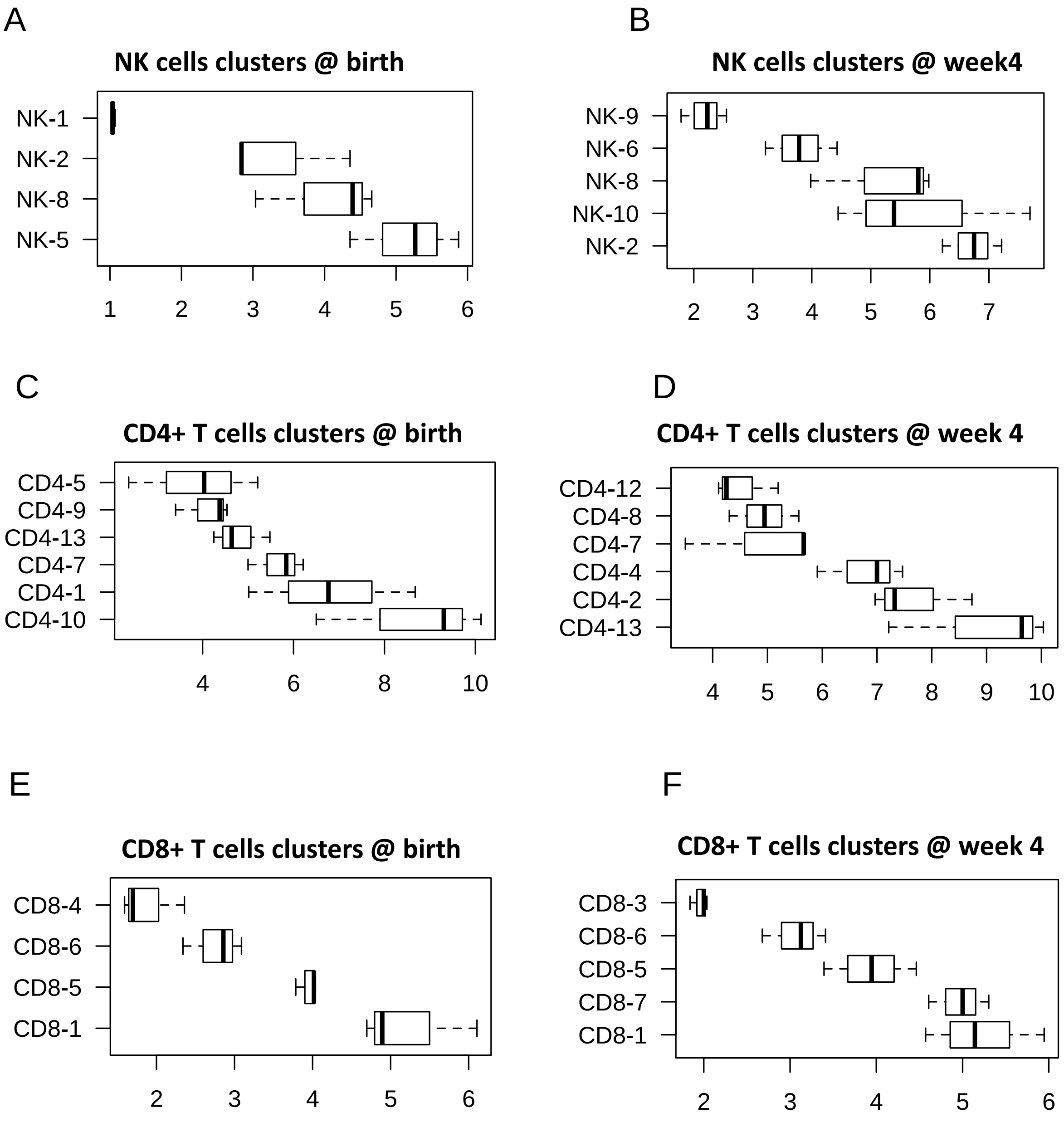

**Figure S6** Immune cell clusters predictive of rotavirus specific antibody responses post-vaccination. Multivariable regression using partial least squares with discriminate analysis (PLS-DA) and recursive variable elimination within repeated double cross-validation for selection of the minimum number of cell clusters with low misclassification error for predicting rotavirus specific IgG responses at week 36. **A & B)** NK cell clusters predictors at birth and week 4 respectively. **C & D) C**D4+ T cell predictors at birth and week 4 respectively. **E & F) C**D8+ T cell predictors at birth and week 4, respectively.
